# Interspecies metabolite transfer and aggregate formation in a co-culture of *Dehalococcoides* and *Sulfurospirillum* dehalogenating tetrachloroethene to ethene

**DOI:** 10.1101/526210

**Authors:** Stefan Kruse, Dominique Türkowsky, Jan Birkigt, Bruna Matturro, Steffi Franke, Nico Jehmlich, Martin von Bergen, Martin Westermann, Simona Rossetti, Ivonne Nijenhuis, Lorenz Adrian, Gabriele Diekert, Tobias Goris

**Affiliations:** Department of Applied and Ecological Microbiology, Institute of Microbiology, Friedrich Schiller University, Jena, Germany; Department Molecular Systems Biology, Helmholtz Centre for Environmental Research – UFZ, Leipzig, Germany; Department of Isotope Biogeochemistry, Helmholtz Centre for Environmental Research – UFZ, Leipzig, Germany; Water Research Institute, IRSA-CNR, Monterotondo, Rome, Italy; Center for Electron Microscopy of the University Hospital Jena, Jena, Germany; Institute of Biochemistry, Faculty of Life Sciences, University of Leipzig, Leipzig, Germany; Chair of Geobiotechnology, Technische Universität Berlin, Germany

**Author notes:** German Institute of Human Nutrition, Department Molecular Toxicology, Research Group Intestinal Microbiology, Potsdam-Rehbrücke (Tobias Goris). Corresponding author: Tobias Goris.

## Abstract

Microbial communities involving dehalogenating bacteria assist in bioremediation of areas contaminated with halocarbons. To understand molecular interactions between dehalogenating bacteria, we co-cultured *Sulfurospirillum multivorans*, dechlorinating tetrachloroethene (PCE) to *cis*-1,2-dichloroethene (*c*DCE), and *Dehalococcoides mccartyi* strains BTF08 or 195, dehalogenating PCE to ethene. The co-cultures were cultivated with lactate as electron donor. In this co-culture, the bacterial cells formed aggregates and *D. mccartyi* established an unusual, barrel-like morphology. An extracellular matrix surrounding bacterial cells in the aggregates enhanced cell-to-cell contact. PCE was dehalogenated to ethene at least three times faster in the co-culture. The dehalogenation was carried out via PceA of *S. multivorans*, and PteA (a recently described PCE dehalogenase) and VcrA of *D. mccartyi* BTF08, as supported by protein abundance. The co-culture was not dependent on exogenous hydrogen and acetate, suggesting a syntrophic relationship in which the obligate hydrogen consumer *D. mccartyi* consumes hydrogen and acetate produced by *S. multivorans*. The cobamide cofactor of the reductive dehalogenase – mandatory for *D. mccartyi* – was also produced by *S. multivorans*. *D. mccartyi* strain 195 dechlorinated *c*DCE in the presence of norpseudo-B_12_ produced by *S. multivorans*, but *D. mccartyi* strain BTF08 depended on an exogenous lower cobamide ligand. This observation is important for bioremediation, since cofactor supply in the environment might be a limiting factor for PCE dehalogenation to ethene, described for *D. mccartyi* exclusively. The findings from this co-culture give new insights into aggregate formation and the physiology of *D. mccartyi* within a bacterial community.

## Introduction

Microbial communities interact in numerous ways, involving the transfer and consumption of metabolic products. Molecular hydrogen, for example, is an important electron carrier in syntrophic communities, in which hydrogen is produced by e.g. fermentation and taken up by hydrogen-consuming prokaryotes. This hydrogen consumption leads to a lower hydrogen partial pressure, which allows otherwise thermodynamically unfavorable reactions to proceed. Therefore, the involved bacteria are physiologically dependent on each other [1, 2, 3]. The probably carcinogenic groundwater pollutant tetrachloroethene (PCE), a remnant of e.g. bleaching, dry-cleaning and metal degreasing, is completely, but often slowly, dechlorinated to ethene in communities involving hydrogen transfer from fermenting bacteria to the obligately hydrogen-consuming *D. mccartyi* [4, 5, 6, 7]. PCE and other organohalides such as hexachlorobenzene or polychlorinated biphenyls are used as terminal electron acceptors by *D. mccartyi* and other bacteria in an anaerobic respiratory chain coupling the dehalogenation to energy conservation via electron transport phosphorylation [6]. This process is termed organohalide respiration (OHR) and involves the corrinoid-containing reductive dehalogenases (RDases) as terminal reductases [8, 9, 10]. However, *D. mccartyi* is characterized by low growth rates and dehalogenates PCE slower [11] compared to other dehalogenating species such as *Sulfurospirillum multivorans. D. mccartyiis* strictly dependent on several specific vitamins, most importantly cobamides [12]. Besides being restricted to hydrogen as electron donor, *D. mccartyi* uses only acetate plus bicarbonate as carbon source and organohalides as electron acceptors. In addition, these bacteria are not able to *de novo* synthesize corrinoids, the obligate cofactor of RDases [11, 13]. While proteins for complete corrinoid biosynthesis are usually not encoded in the genomes of *D. mccartyi* [14], different studies revealed its ability to salvage and remodel corrinoids, enabling *D. mccartyi* to restore dechlorination [15, 16, 17]. The functionality of the corrinoid and thus the RDase is directly dependent on the type of the lower base in *D. mccartyi*. Only three types of corrinoids have so far been described to be functional in *D. mccartyi* strain 195: 5,6-dimethylbenzimidazolyl-cobamide ([DMB]-Cba), 5-methylbenzimidazolyl-cobamide and 5-methoxybenzimidazolyl-cobamide ([5-OMeBza]Cba). Nonfunctional corrinoids e.g. 5-hydroxybenzimidazolyl-cobamide or 7-adeninyl-cobamide ([Ade]Cba) can be converted to functional ones by replacement of the lower ligand in *D. mccartyi* when 5,6-dimethylbenzimidazole (DMB) is provided [16, 17].

Several studies on dechlorinating communities containing *D. mccartyi* in association with fermenting, acetogenic and/or methanogenic bacteria revealed higher dechlorination and growth rates than those of pure cultures [16, 18, 19, 20, 21, 22, 23, 24, 25, 26]. It is hypothesized that cross-feeding and a constant supply of growth factors such as corrinoids and biotin enhance growth [27]. Within these communities, non-dechlorinating fermenting bacteria provide hydrogen, acetate and CO2 from e.g. lactate or butyrate fermentation. The fermenting bacteria are dependent on hydrogen consumers which keep the hydrogen partial pressure low [18, 21, 22, 28]]. For example, co-culture experiments revealed an interspecies hydrogen transfer between *Desulfovibrio desulfuricans* fermenting lactate and *D. mccartyi* dechlorinating TCE [15]. Acetogens like *Sporomusa ovata* and sulfidogens (*Desulfovibrio*), produce different types of corrinoids which can be used by *D. mccartyi* [29, 30]. An interspecies cobamide transfer was also shown between *Methanosarcina barkeri* strain Fusaro and *D. mccartyi* strain BAV1, GT and FL2, when DMB was present [25, 26]. In the only co-culture of *D. mccartyi* (strains BAV1 and FL2) with a PCE-dechlorinating bacterium, *Geobacter lovleyi*, a corrinoid transfer was also observed [31]. In this co-culture, however, hydrogen had to be supplemented. All these studies showed more robust growth of *D. mccartyi* in co-cultures, resulting in higher dechlorination rates and cell yields than in pure cultures and sometimes, depending on the cobamide metabolism of the co-culture, a different degree of the dechlorination extent [32]. However, a PCE-dechlorinating bacterium providing hydrogen, acetate and corrinoids to *D. mccartyi*, to our knowledge, has not been described in the literature. Thus, further information on *D. mccartyi* co-cultures is needed to gain insights into complex microbial communities involving *D. mccartyi*. Relatively simple co-cultures provide a model community in which metabolic transfers are easier to follow. Recently, the PCE to *c*DCE-respiring *Sulfurospirillum multivorans*, capable of *de novo* corrinoid production [33, 34, 35], was shown to produce hydrogen and acetate under fermentative growth conditions, e.g. with pyruvate or lactate and without electron acceptor present [36]. In the same study, a co-culture of *S. multivorans* with a methanogen dependent on hydrogen as electron donor was established. Therefore, *S. multivorans* is a promising partner for a co-cultivation with *D. mccartyi*. In addition, it is of interest, whether *D. mccartyi* is able to take up and utilize norpseudo-B_12_ ([Ade]NCba), so far known to be produced exclusively by *S. multivorans* and *S. halorespirans* [35, 37].In this study, we investigated the physiological interaction between *D. mccartyi* strain BTF08 and 195 in co-culture with *S. multivorans* dechlorinating PCE to ethene by using dechlorination analytics, microscopy and proteomics.

## Materials and Methods

### Cultivation of pure cultures

*D. mccartyi* cultures BTF08 and 195 (maintained at the UFZ Leipzig) were cultivated in 200 mL serum bottles containing 100 mL bicarbonate-buffered mineral salt medium with 5 mM acetate and 148 nM vitamin B_12_ (cyanocobalamin, ca. 200 μg/L), reduced with Na_2_S · 7 H_2_O (final concentration, 0.25 g/L) [38]. Anoxic atmosphere was established by 30 cycles of gassing and degassing with nitrogen, and CO_2_ was added to a final atmosphere of N_2_:CO_2_ (75:25 v/v 150 kPa). After autoclaving the medium and adjusting the pressure to approximately 100 kPa with nitrogen and CO_2_, we applied, in addition, hydrogen [50 kPa] to avoid possible electron donor limitation. PCE (>99% purity, Sigma Aldrich, Steinheim, Germany) and *c*DCE (97% purity, Sigma Aldrich, Steinheim, Germany) served as electron acceptors and were added with a microliter syringe (Hamilton, Bonaduz, Switzerland) to a final concentration of 0.35 mM (aqueous phase concentration). Re-feeding of the cultures was done with the same amount of PCE or *c*DCE after complete conversion to ethene. After maximally three re-feeding steps, cultures were transferred [10% (v/v)] into fresh medium. To evaluate the effect of different types and concentrations of B_12_ on dechlorination activities, *D. mccartyi* cultures received 54 nM norpseudo-B_12_ ([Ade]Cba) or 5-OMeBza-B_12_ ([5-OMeBza]Cba), each. Norpseudo-B_12_ was extracted as described previously [39] from 6 L of *S. multivorans* grown anaerobically with 40 mM pyruvate and 10 mM PCE as described elsewhere [40]. In brief, crude cell extract was acidified to a pH of ≤ 5 with acetic acid before adding 0.1 M potassium cyanide and boiling the sample for 15 min. Afterwards, the sample was centrifuged and the supernatant kept on ice. The cell pellet was reuspendend in 10 mL UPW and the same extraction procedure was applied twice. The combined supernatant was mixed with Amberlite XAD4 (0.25 g XAD4 per ml extract; Sigma-Aldrich) and incubated on a shaker overnight. The supernatant was removed and the cobamide-loaded XAD4 material was washed with 10 volumes of UPW. Elution of cobamide was done for 1 h on a shaker with 1 volume of methanol which was repeated twice. A vacuum concentrator was used to completely dry the eluates which were then resuspended in 2 mL UPW. The sample was applied on a column containing 3 g aluminium oxide, recovered with 50 mL UPW and dried. Resuspension of purified Norpseudo-B_12_ was done with UPW. 5-OMeBza-B_12_ was obtained in the same way from 6L of *Desulfitobacterium hafniense* DCB2 grown anaerobically with 40 mM pyruvate and 10 mM ClOHPA (3-chloro-4-hydroxy-phenylacetate) on a medium amended with 25 μM 5-OMeBza [41]. *S. multivorans* (DSMZ 12446) was cultured in the same mineral salt medium as *D. mccartyi* [38] with 40 mM lactate and 10mM nominal PCE in hexadecane (2 mL PCE/hexadecane per 100 mL medium) but without the addition of acetate and hydrogen [38]. All cultivation experiments were performed statically at 28°C in the dark and in biological triplicates.

### Co-cultivation of *S. multivorans* and *D. mccartyi*

Co-cultures of *S. multivorans* and *D. mccartyi* strain BTF08 or 195 were maintained in the same mineral salt medium as the pure cultures, without acetate and hydrogen. Initially, the inoculum of the pure cultures into the co-culture was 10 % of each pure culture. The medium contained 25 mM lactate (D/L, > 85% purity, Sigma-Aldrich, Steinheim, Germany) as the electron donor for *S. multivorans* and 0.35 mM PCE (aqueous-phase concentration) as the electron acceptor. Lactate (25 mM) and PCE (0.35 mM) was re-fed after depletion of substrates was measured. The B_12_-dependance of the co-cultures was tested with 148 nM vitamin B_12_ (>98% purity, Sigma-Aldrich, Steinheim, Germany) serving as the positive control and without vitamin B_12_ amendments. Cultures without vitamin B_12_ received 1 μM DMB (>99% purity, Sigma-Aldrich, Steinheim, Germany), where indicated. The DMB dose used had only minimal negative effect on *S. multivorans* growth [39]. For isotope fractionation experiments, *S. multivorans* and *Sm*/BTF08 co-cultures were cultivated in 50 mL serum bottles with 25 mL medium. Replicate bottles (triplicates for each time point) were inoculated at the same time and the dehalogenation process was stopped at different time points by addition of 3 mL 2 M Na_2_SO_4_ (pH 1.0).

### Quantitative (q)PCR analysis of cell growth

Quantitative PCR (qPCR) was applied to count *Sulfurospirillum* and *Dehalococcoides* 16S rRNA gene copies. DNA was extracted from 1 mL co-culture taken from different time points during the cultivation experiment using the NucleoSpin Tissue DNA extraction kit according to the manufacturer’s instructions (Macherey-Nagel, Düren, Germany). Cell numbers were quantified by direct epifluorescence microscopy of Sybr-green stained cells immobilized on agarose-coated slides to generate standard curves for *q*PCR as described previously [42]. For the corresponding standard curves, *D. mccartyi* and *S. multivorans* cell suspensions with known cell numbers (1×10^4^ to 1×10^8^ and 1×10^5^ to 1×10^9^ respectively) were subjected to qPCR. The qPCR reaction mixture contained 1 μl of gDNA or DNA standard, 6.25 μl 1x KAPA SYBR Fast master mix (Sigma Aldrich, Steinheim, Germany) and 0.208 μM forward and reverse primer. Primers used were Dhc_sp_16S_fw (5’-GTATCGACCCTCTCTGTGCCG-3’) and Dhc_sp_16S_rev (5’-GCAAGTTCCTGACTTAACAGGTCGT-3’) for *D. mccartyi* and Smul_16S_fw (5’-AGGCTAGTTTACTAGAACTTAGAG-3’) and Smul_16S_rev (5’-CAGTCTGATTAGAGTGCTCAG-3’) for *S. multivorans*. The conditions of the PCR program were as followed: 95°C for 2 min (initial denaturation) followed by 40 cycles of 55°C (*S. multivorans* primer) or 60°C (*D. mccartyi* primer) for 20s (annealing), 72°C for 30s (elongation) and 95°C for 10 s (denaturation). Each *q*PCR program included a melting curve for verification of specific target DNA amplification and primer efficiency was tested for used primers (R^2^ *Sm*: 99.7%; R^2^ *Dhc*: 98.8%, see Supplementary Figure S1). The cell numbers of samples were calculated by comparing the sample CT values with the CT values of the standard curve obtained from microscopical cell counting (Supplementary Figure S1). All samples were conducted in three biological replicates with two corresponding technical replicates and three technical replicates for the calibration curve.

### Analytical methods

Ethene and chlorinated ethenes were quantified gas-chromatographically with a flame ionization detector (Clarus 500, Perkin Elmer, Rodgau, Germany) and a CP-PoraBOND Q FUSED SILICA 25 m x 0.32 mm column (Agilent Technologies, Böblingen, Germany). Samples from the cultures (1 mL of the gas phase and 1 mL of the liquid phase) were transferred to GC vials with gast tight syringes, the sampling proceeded via a headspace sampler (HS 40, Perkin Elmer; needle temperature 100°C, vial pressurized with 0.1 kPa nitrogen for 2 mins). The GC vial with the liquid phase was heated in the HS 40 to 80°C for 5 min. Samples were transferred through the transfer line (100°C) into the GC. Ethene and chlorinated ethenes were separated with 4 min at 60°C rising to 280°C in 10°C min^-1^ steps. The injector temperature was fixed at 250°C and the detector temperature at 300°C. Standard curves of ethene and each chlorinated ethene were recorded for peak area quantification and retention times were compared to known standards. Nonane served as an internal standard during measurements. The retention times were: PCE (detection limit [dl] 2.1 μM), 15.1 min; TCE (dl 2.5 μM) 13.2 min; *c*DCE (dl 3.1 μM), 11.3 min; VC (dl 3 μM), 6.0 min; ethene (dl 2.5 μM), 1.3 min. Due to the low solubility of PCE, it was added to the medium followed by stirring for at least 8 hours to allow it to completely dissolve. This lead to a reliable quantification of only the initial PCE concentration, but not of the re-fed PCE, especially since a portion of the re-fed PCE is rapidly degraded by the now higher abundant *S. multivorans* cells. The concentrations of chlorinated ethenes (in μM or mM) were calculated by summing up the concentrations in the gas phase and liquid phase. Dechlorination rates were calculated according to the average amount of stoichometrically converted PCE to the corresponding end products VC (in the case of *D. mccartyi* strain 195, where the formation of co-metabolically produced ethene was negligible) or ethene (for *D. mccartyi* strain BTF08) divided by time in days. For example the conversion of 400 μM PCE into 400 μM ethene over 10 days resulted in a dechlorination rate of 40 μM PCE/day. Hydrogen (dl 0.8 μM) was measured gas-chromatographically using a thermal conductivity detector (AutoSystem, Perkin Elmer, Rodgau, Germany) equipped with a Carboxen-1010 PLOT capillary column 30 m x 0.53 mm (Supelco, München, Germany) column. Separation of analytes was done at a temperature of 35°C (detector at 150°C). For this, the culture headspace (1 mL) was sampled with gas-tight syringes (Hamilton, Bonaduz, Switzerland) and injected directly into the injection valve of the GC. Concentrations were calculated using calibration curves ranging from 4 – 50 μmol hydrogen. Organic acids (lactate, acetate, pyruvate and succinate) were analyzed using a reversed-phased HPLC (Merck-Hitachi, Darmstadt, Germany) equipped with a diode array detector (L-7450) and separated on an AMINEX HPX-87H column (7.8 x 300 mm; BioRad, Munich, Germany) with a cation H guard pre-column. Samples (1 mL) from the liquid phase of the cultures were filtered (MiniSart RC4, 0.2 μm, Sartorius, Göttingen, Germany) and acidified with 2.5 μL H_2_SO_4_. Of this sample, 20 μL was injected into the HPLC. Monitoring of organic acids was based on their absorption at 210 nm. The column was operated at a temperature of 50°C and 5 mM (v/v) H_2_SO_4_ served as mobile phase with a flow rate of 0.7 mL min^-1^. The retention times were: lactate (dl 0.2 mM), 10.61 min; pyruvate (dl 0.01 mM) 7.92 min; acetate (dl 0.4 mM), 12.88 min; succinate (dl 0.15 mM), 9.81 min. Peak areas and retention times were related to standards ranging from 1 to 10 mM or 0.5 to 2 mM (for pyruvate).

### Compound-specific stable isotope analysis

Determination of the carbon isotope composition of the chlorinated ethenes in pure culture of *S. multivorans* and in co-culture of *S. multivorans* and *D. mccartyi* strain BTF08 was done using gas chromatography combustion isotope ratio mass spectrometry (GC-C-IRMS; Thermo GC Trace 1320 combined with Thermo-Finnigan MAT 253 IRMS, Bremen, Germany) [43]. All samples were analyzed in technical triplicates. 2 mL liquid phase were taken from the respective sample and transferred to a He-flushed 10 mL crimped vial. Of these, 1 mL was taken from the headspace via an autosampler (Thermo TriPlus RSH Autosampler, Bremen, Germany) and injected in a gas chromatograph with a split ratio of 1:5. Chlorinated ethenes were separated on a DB-MTBE column (60 m x 0.32 mm x 1.8 μm, J&W Scientific, Waldbronn, Germany) and the following temperature program: 40°C for 5 min, increase to 250°C by 20°C min^-1^ and hold for 5 min using helium as carrier gas at a flow rate of 2.0 mL min^-1^ (Injector at 250°C).

The carbon isotope composition is given in the δ-notation (‰) relative to the Vienna Pee Dee Belemnite standard [44]. Carbon isotope fractionation was calculated using the Rayleigh equation (eq 1) where R_0_ and R_t_ represent the isotope values and C_0_ and C_t_ the concentrations at time 0 and t [45, 46].

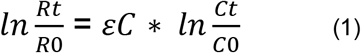

The carbon isotope enrichment factor (ε_C_) relates changes in the concentration of the isotopes to changes in their isotope composition. A two-tailed T-Test was used to calculate the 95 % confidence interval based on the slope. Standard deviations were obtained from at least triplicate measurements (< 0.5 ‰).

### B_12_ extraction and MS analysis

The B_12_ content of three independently cultivated 100 mL cultures was analyzed. For this, the culture volume was reduced to 20 mL using a vacuum concentrator and 0.1 M potassium cyanide was added. After boiling the samples for 20 min, cell debris was removed by centrifugation (10 min, 6700 x g, 8°C). The supernatant was applied onto a C-18 column (CHROMABOND C-18 ec, Macherey-Nagel, Düren, Germany) equilibrated with 5 mL 100% (v/v) methanol and 5 mL ultrapure water (UPW). Washing of the column was done twice with 5 mL UPW and B_12_-types were eluted with 5 mL 100% (v/v) methanol. The eluate was completely dried in a vacuum dryer and dissolved in UPW prior to analysis. The extract was injected to ultrahigh performance nano-flow liquid chromatography (UHPLC) (Ultimate 3000, Thermo Fisher, Waltham, USA) coupled to mass spectrometer (LC-MS, Orbitrap Fusion, Thermo Fisher, Waltham, USA) via heated electrospray ionization (HESI-II, Thermo Fisher, Waltham, USA). The UHPLC was equipped with a Hypersil Gold C18 column (150 x 2.1 mm, 3 μm film thickness; Thermo Fisher, Waltham, USA) and a C18 guard column (10 x 2.1 mm, Waters, Milford, USA). Chromatographic separation using a gradient method with 0.1% formic acid (A) and methanol (B) as mobile phase was applied as following: 5% B for 1 min, 60 min gradient to 90% B, 4 min at 90% B, 1 min gradient to 5% B and 4 min at 5% B with a constant flow of 0.2 mL min^-1^ and 25°C column oven temperature. Injection volume was 5 μL. Ionization was set to positive ion mode at 3.5 kV, 35 arbitrary units (Arb) sheath gas, 10 Arb auxillary gas, 325°C ion transfer tube temperature and 275°C vaporizer temperature. Orbitrap resolution for precursor scan (MS1) was set to 120.000 with a scan range of 300-1600 *m/z*. Data evaluation was done on the original mass spectra comparing with predicted masses of corrinoids (Supplementary Tables S1 and S2, for cobalamin standards see Supplementary Figure S2).

### Scanning Electron Microscopy and fluorescence in situ hybridization

Field emission-scanning electron microscopy (FE-SEM) was performed with co-cultures of *S. multivorans* and *D. mccartyi* strains BTF08 and 195 and pure *S. multivorans* and *D. mccartyi* strain BTF08 cultures. All of the cultures were treated the same way for FE-SEM as follows. Cells of 5 mL culture were incubated for 15 min with 2.5% glutaraldehyde and pre-fixed for 2 h on poly-L-lysine coated coverslides (12 mm, Fisher Scientific, Schwerte, Germany). Cover slides were washed three times with 0.1 M sodium cacodylate (pH 7.2) (>98% purity, Sigma Aldrich, Steinheim, Germany) and post-fixed for 1 h with 1% osmium tetroxyde in the same cacodylate buffer. After fixation, samples were dehydrated using different ethanol concentrations (35%/50%/70%/80%/95%/100% v/v) and incubated for 10 min at each step. Critical point drying was done in a Leica EM CPD200 Automated Critical Point Dryer (Leica, Wetzlar, Germany), followed by coating with 6 nm platinum in a BAL-TEC MED 020 Sputter Coating System (BAL-TEC, Balzers, Liechtenstein). Imaging of the samples was done with a Zeiss-LEO 1530 Gemini field emission scanning electron microscope (Carl Zeiss, Oberkochen, Germany) at different magnifications and at 10 kV acceleration voltage. Fluorescence in situ hybridization (FISH) was performed as described previously [47, 48]. In brief, samples were fixed with formaldehyde (2% v/v final concentration) for 2 h at 4°C and filtered on polycarbonate membrane filters (47 mm diameter, 0.2 μm pore size, Nucleopore). Filters were stored at −20°C until further processing. FISH detection of *D. mccartyi* strain BTF08 and 195 was done with Cy3-labeled DHC1259t and DHC1259c probes and *S. multivorans* detection with FITC-labeled probes SULF F220ab [49]. Imaging of un-aggregated cells was done with an epifluorescence microscope (Olympus, BX51) combined with an Olympus XM10 camera. Images were analyzed via Cell-F software. Aggregates were visualized using a confocal laser scanning microscopy (CSLM, Olympus FV1000).

### Protein extraction and proteome analysis

Samples for proteome analyses were taken approximately two weeks after one re-feeding with PCE or *c*DCE (pure *D. mccartyi* strain BTF08 cultures) or 12 hours after the third re-feeding with PCE (co-cultures *Sm*/BTF08, Supplementary Figure S3). When harvested, PCE, TCE and *c*DCE were the dominant chlorinated ethenes in the pure cultures, while *c*DCE and VC were the dominant chlorinated ethenes in the co-cultures (Supplementary Figure S3). Samples were processed as described previously [50]. Briefly, protein extraction was performed in lysis buffer (20 mM HEPES, 1 mM sodium vanadate, 1 mM β-glycerol phosphate, 2.5 mM sodium pyrophosphate, 8 M urea [51]) by three freeze/thaw-cycles and ultrasonic bath treatments. Ten μg protein dissolved in 100 μL ultra pure water was precipitated with the five-fold amount (v/v) of ice-cold (−20°C) acetone. Protein pellets were dissolved in 50 μL SDS sample buffer (2% SDS, 2 mM beta-mercaptoethanol, 4% glycerol, 40 mM Tris-HCl pH 6.8, 0.01% bromophenolblue), heated to 90 °C for 5 min and separated on a SDS gel (12.5%). The gel was run until the samples entered the separating gel. Afterwards, a 3-5 mm protein band from each sample was cut out, destained, dehydrated, reduced with 10 mM dithiothreitol, alkylated with 100 mM iodoacetamide and proteolytically cleaved over night at 37°C using trypsin (Promega, Madison, WI, USA). Peptides were extracted, desalted using C18 ZipTip columns (Merck Millipore, Darmstadt, Germany) and resuspended in 0.1% (v/v) formic acid before LC–MS analysis.

Proteolytic lysates were separated using an Ultimate 3000 RSLCnano liquid chromatographic instrument (Thermo Scientific, Germany). Mass spectrometry was performed on an Orbitrap Fusion mass spectrometer (Thermo Scientific, San Jose, CA, USA) coupled to a TriVersa NanoMate (Advion, Ltd., Harlow, UK). Samples of 5 μL were loaded onto a trapping column (Acclaim PepMap100 C18, 75 μm × 2 cm, Thermo Scientific) using 96% eluent A (0.1% formic acid) and 4% eluent B (0.08% formic acid, 80% acetonitrile) at a flow rate of 5 μL min^-1^ and separated via a 25 cm analytical column (Acclaim PepMap100 C18, 75 μm × 25 cm, Thermo Scientific) at 35°C using a constant flow rate of 300 nL/min. Peptide separation was achieved by applying a linear gradient of eluent B from 4% to 50% within 100 min. Full MS scans were measured in the Orbitrap mass analyzer within the mass range of 400-1,600 *m/z* at 120,000 resolution using an automatic gain control (AGC) target of 4×10^5^ and maximum fill time of 60 ms. The MS instrument measured in data-dependent acquisition (DDA) mode using the highest intense ion (top speed, 3 sec cycle time). Positive ion charge states between 2 and 7 were selected for MS/MS. Precursor masses for MS/MS were selected based on the highest intensity and excluded from further MS/MS for 30 s to prevent redundancy in MS/MS acquisition. After higher energy collisional (HCD) fragmentation at normalized collision induced energy of 30%, fragment masses were scanned in the Orbitrap mass analyzer at a resolution of 15,000 with 5×10^4^ AGC target and a maximum injection time of 150 ms.

LC-MS/MS data were analyzed using Proteome Discoverer (v2.1, Thermo Scientific). MS/MS spectra were searched against a combination of a *S. multivorans* (3,233 non-redundant protein-coding sequences, downloaded January 2017 from NCBI GenBank, accession number CP007201.1) and a *D. mccartyi* strain BTF08 database (1,535 non-redundant protein-coding sequences, downloaded December 2016 from NCBI, accession number CP004080.1). A “common repository of adventitious proteins database” (cRAP) was integrated to ensure correct protein identifications. The SEQUEST HT algorithm was used with the following settings: trypsin as cleavage enzyme, oxidation on methionine as dynamic and carbamidomethylation on cysteine as static modification, up to two missed cleavages, precursor mass tolerance set to 10 ppm and fragment mass tolerance to 0.02 Da, respectively. Only peptides with a false discovery rate (FDR) 1%, XCorr ≥2, q-value and the posterior error probability (PEP) ≤0.01 were considered as identified (Supplementary Excel File S1). Quantification of proteins was performed using the average of the top three peptide MS1 areas. After log10 transformation, the protein values were normalized to the median of all proteins of a sample, to the median of all proteins of the respective organism in that sample and the median of all samples and scaled so that the global minimum is zero. Two outliers of the co-culture proteome replicates with only 11 (C3) and 92 (P1, see Supplementary Excel File S1) protein identifications, compared to at least 317 protein identifications from the other *D. mccartyi* pure cultures, were excluded from the analysis. Proteins with only one out of three possible quantitative values per sample were considered as identified only. P-values were calculated using a two-tailed, homoscedastic student’s t-test and multiple corrected with the Benjamini-Hochberg method. Figures were created using an in-house written R-script with the packages gplots, ggplot2, ggbiplot, dplyr, miscTools and vegan. The non-parametrical multiple dimensional scaling (nMDS)-analysis was performed with the anosim-function of the vegan package in R (v3.4.1). *Pairwise Indicator Species Analysis* was used to identify proteins that were significantly associated with the different cultivation conditions [52, 53]. Indval scores and significances were calculated according to Malik et al. [54].

## Results

### 1. Growth and dechlorination in *Sulfurospirillum* (*Sm*)/*Dehalococcoides* (*Dhc*) co-cultures

The growth characteristics and PCE to *c*DCE dechlorination pattern of *S. multivorans* cultivated in the co-culture medium (Supplementary Figure S4) was comparable to those in the routinely used medium [55]. With lactate as electron donor and PCE as electron acceptor, a co-culture of *S. multivorans* and *D. mccartyi* BTF08 maintained over 10 transfers under the same conditions dechlorinated PCE to stoichiometric amounts of ethene within eight days after inoculation (Figure 1A). The dechlorination rate (PCE to ethene) of the *Sm*/BTF08 co-culture was 4.5-fold higher compared to the *D. mccartyi* strain BTF08 pure culture (Table 1). In the latter, PCE was completely reduced to ethene within 35 days at a rate of 9 ± 0.3 μM/day (Supplementary Figure S5A). The culture was re-fed with PCE at day 8 and day 12 after inoculation. After re-feeding, the dechlorination increased (Figure 1A) and complete dechlorination of PCE occurred within three days. PCE was dechlorinated to *c*DCE within two days. To check whether *S. multivorans*, known for its high dehalogenation rate [55], was responsible for this fast dechlorination, the stable carbon isotope fractionation patterns (which largely differ between *D. mccartyi* and *S. multivorans*) of pure and co-cultures were compared. The isotope fractionation pattern of PCE in the co-culture did not change significantly, and ranged from −30.2 ± 0.09 to −29.5 ± 0.12 ‰ at 70% of transformed PCE (Supplementary Figure S6A). A low but significant fractionation was measured in the pure culture of *S. multivorans* (from −29.2 ± 0.05 to −27.6 ± 0.11 ‰ at 56% of transformed PCE, Supplementary Figure S6A). The isotope enrichment factors for the *S. multivorans* pure and the co-culture were in the same range while differing largely from that of a *D. mccartyi* strain BTF08 pure culture (*Sm*/BTF08 co-culture: ε_C_ = −0.4 ± 0.3 ‰; *S. multivorans* pure culture: ε_C_ = −2.0 ± 0.4 ‰, Supplementary Figure S6B; *D. mccartyi* strain BTF08: ε_C_ = −5.0 to −9.0 ‰ [56]). This high similarity in the PCE isotope fractionation patterns of *S. multivorans* pure and co-cultures is likely due to PCE to cDCE dechlorination performed nearly exclusively by *S. multivorans*. The fast dechlorination of PCE to *c*DCE in less than two days was also reflected in the increase of the *S. multivorans* cell number (at least one cell doubling per day), until PCE was completely dechlorinated to *c*DCE (Figure 1C). After this respiratory growth, a weaker growth (less than one cell doubling in 20 days) was observed, probably correlating to lactate fermentation of *S. multivorans* (Figures 1E, 1F). *D. mccartyi* strain BTF08 needed seven days for one cell doubling in the co-culture. For both organisms, a correlation between dechlorination and growth was observed. The ratio between *S. multivorans* and *D. mccartyi* strain BTF08 cells changed from initially 2.6 to 5.9 after 2 days and 4.6 after 15 days and two re-feeding steps. After 12 days, when the second dose of PCE was completely dechlorinated to ethene, lactate was completely consumed, therefore lactate and PCE were re-fed. Acetate production occurred continuously during the whole dechlorination process (Figure 1E), other organic acids such as succinate were not detected. No hydrogen was detected in the gas phase at any point. Because the concentration of hydrogen and succinate which were shown to be produced during pyruvate fermentation of *S. multivorans* pure cultures [36] might be below the detection limit in our setup and pure *S. multivorans* cultures could not grow via lactate fermentation, a stoichiometry could not be calculated.

**Table 1:** Dechlorination rates of different co-culture set-ups *of S. multivorans* and *D. mccartyi* BTF08 or 195 from PCE to the products VC (*D. mccartyi* strain 195) or ethene (*D. mccartyi* strain BTF08).

| Organism(s) | Electron donor | Dechlorination rate ($\mu\text{M day}^{-1}$) |
| --- | --- | --- |
| **With vitamin B<sub>12</sub>** |  |  |
| *D. mccartyi* BTF08 | H<sub>2</sub> | 9 ± 0.3 |
| *D. mccartyi* 195 | H<sub>2</sub> | 15 ± 0.4 |
| *Sm*/BTF08 | Lactate | 41 ± 2 |
| *Sm*/195 | Lactate | 48 ± 1 |
| **Without vitamin B<sub>12</sub>** |  |  |
| *Sm*/BTF08 | Lactate | - |
| *Sm*/195 | Lactate | 9 ± 0.04 |
| **Without vitamin B<sub>12</sub> / with DMB** |  |  |
| *Sm*/BTF08 | Lactate | 36 ± 2 |
| *Sm*/195 | Lactate | 8 ± 0.5 |

∴ No dechlorination to VC or ethene observed.

**Figure 1:**
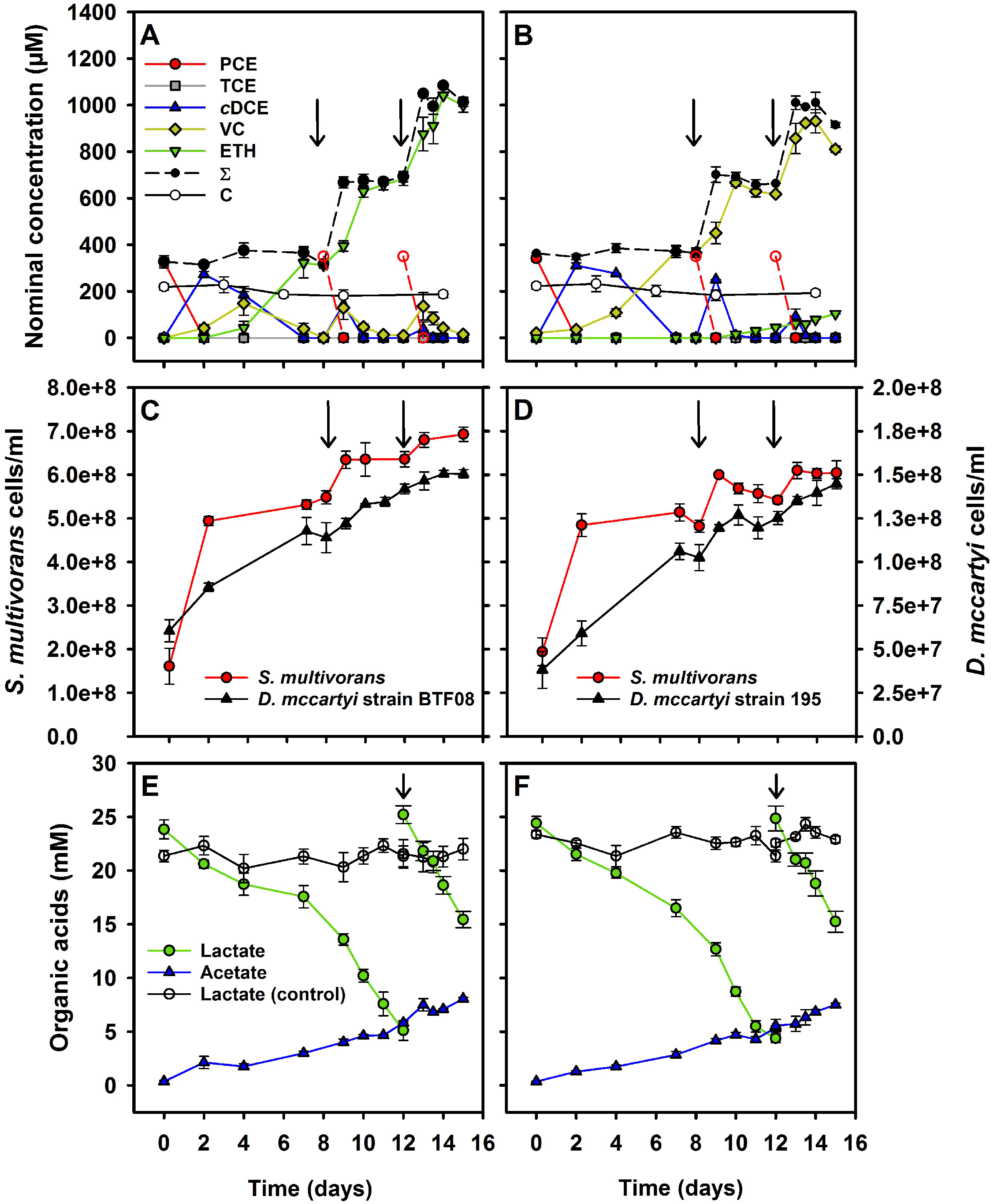
Dechlorination of chlorinated ethenes, growth and metabolite analysis of *S. multivorans*/*D. mccartyi* co-cultures with vitamin B_12_ amendment. (A) PCE dechlorination of *Sm*/BTF08 and (B) *Sm*/195. (C) Growth curve of *Sm*/BTF08 and (D) *Sm*/195. (E) Lactate consumption and acetate production of *Sm*/BTF08 and (F) *Sm*/195. Arrows indicate the time points of re-feeding the culture with PCE (A-D) or lactate (E and F). Broken red lines with open symbols represent theoretical, not analytical, values of the PCE concentration as added to the culture. Please note the secondary y-axes for *D. mccartyi* cell numbers in C and D. Negative controls were run with autoclaved cells (abiotic controls, C). Standard deviation of three independent biological replicates (N=3) are represented by error bars (not visible when smaller than the used symbol). *∑* – mass balance; sum of PCE, TCE, *cis*-DCE, VC and ethene.

Similar growth characteristics and dechlorination behavior were observed in the *Sm*/195 co-culture cultivated under the same conditions, except that vinyl chloride (VC) was the major dechlorination product. The first dose of PCE was dechlorinated within 7 days stoichiometrically to VC (Figure 1B). The increase of cell number was slightly lower for *S. multivorans* and slightly higher for *D. mccartyi* strain 195 compared to the *Sm*/BTF08 co-culture (Figure 1D). After the second dose of PCE was dechlorinated to VC, a low amount of ethene was produced starting on day 10, reaching 104 μM ethene after day 15. The *Sm*/195 co-culture reduced PCE to VC more than 3-fold faster than the *D. mccartyi* strain 195 pure culture, which completed dechlorination to VC within 18 days (Table 1, Supplementary Figure S5B).

### 2. Corrinoid transfer in co-cultures and the effect of the lower corrinoid ligand

*D. mccartyi* strains rely on externally provided corrinoids for dehalogenation and growth. This was also tested for *D. mccartyi* strains BTF08 and 195, which only showed negligible *c*DCE dechlorination after 100 days without addition of corrinoid (Figure 2A and D). Therefore, it was of interest whether *S. multivorans* is able to provide functional corrinoids for *D. mccartyi* strains BTF08 and 195. In *Sm*/BTF08 co-cultures without the amendment of vitamin B_12_, stoichiometric dechlorination of PCE to *c*DCE was obtained (~30 μmol/bottle/day) (Figure 2B). No further dechlorination of *c*DCE to VC or ethene was detected, indicating that *S. multivorans* alone was responsible for the dechlorination. When *c*DCE dechlorination stalled in the *Sm*/BTF08 co-culture, 1 μM 5,6-dimethylbenzimidazole (DMB) was added, which resulted in subsequent *c*DCE dechlorination (Figure 2C). Ethene production rates from *c*DCE (36 ± 2 μM/day) were similar to the co-culture amended with vitamin B_12_ (41 ± 2 μM/day). The dechlorination rate increased after re-feeding with PCE to 8 μmol/bottle/day. In the *Sm*/BTF08 co-culture with DMB but without vitamin B_12_ amendment, three different types of corrinoids were found by mass spectrometric analysis: [Ade]NCba, [DMB]NCba and [DMB]Cba (Supplementary Figure S7).

**Figure 2:**
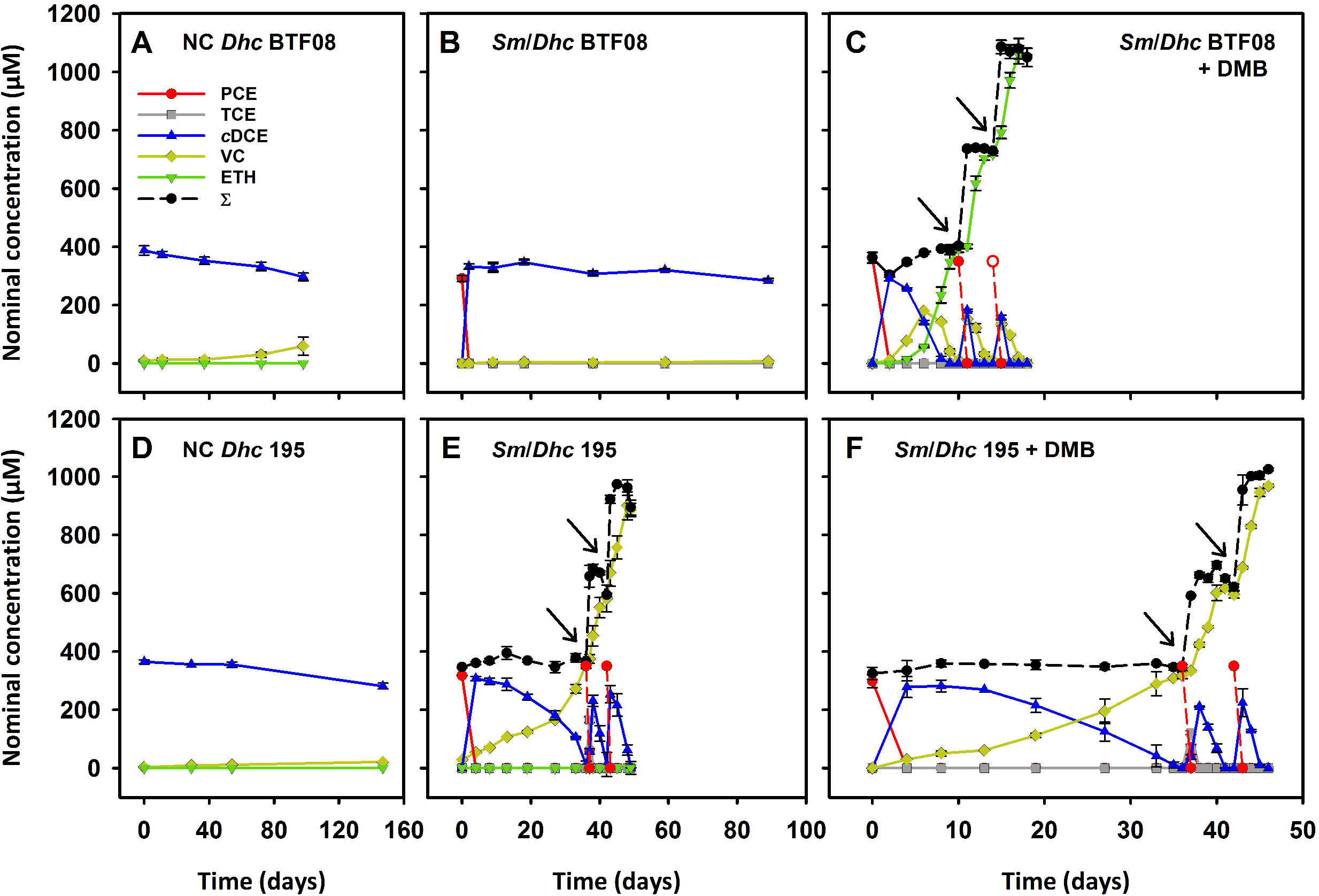
Dechlorination of *S. multivorans/D. mccartyi* co-cultures without addition of vitamin B_12_. (A) Strain BTF08 with *c*DCE as electron acceptor (negative control). (B) *Sm*/BTF08 with PCE as the electron acceptor. (C) *Sm*/BTF08 with PCE as electron acceptor and amendment of 1 μM DMB. (D) Strain 195 with *c*DCE as electron acceptor (negative control). (E) *Sm*/195 with PCE as the electron acceptor. (F) *Sm*/195 with PCE as electron acceptor and amendment of 1 μM DMB. Please note the different time scales. All growth experiments were conducted in biological triplicates (N=3). Arrows indicate re-feeding of PCE. ∑ – mass balance; sum of PCE, TCE, *c*DCE, VC and ethene.

Interestingly and in contrast to the *Sm*/BTF08 co-culture, the *Sm*/195 co-culture without vitamin B_12_ amendment (-B_12_) dechlorinated PCE to VC, although at low rates (Figure 2E). After 35 days, PCE was re-fed, and the *c*DCE to VC dechlorination rate increased 4-fold (3.6 μmol/bottle/day). The only corrinoid detected in the *Sm*/195 co-culture-B_12_ was [Ade]NCba (Supplementary Figure S8). No significant increase in *c*DCE dechlorination was observed when 1 μM DMB was added to the *Sm*/195 co-culture (Figure 2F). Similar to the *Sm*/BTF08 co-culture, [Ade]NCba, [DMB]NCba and [DMB]Cba were found by mass spectrometric analysis when DMB was added to the *Sm*/195 co-culture (Supplementary Figure S9). To confirm that *D. mccartyi* strain 195 can use [Ade]NCba for dechlorination of *c*DCE to VC, we amended a pure culture with [Ade]NCba isolated from *S. multivorans* (Supplementary Figure S10A). Dechlorination of 50 μmol *c*DCE to VC was completed within 22 days (Supplementary Figure S10A). *D. mccartyi* strain 195 pure cultures amended with [5-OMeBza]Cba showed slightly faster dechlorination and the *c*DCE was converted within 16 days into VC (Supplementary Figure S10B). Only the amended corrinoid types were detected in the cultures via MS, impurities and rearrangement of corrinoids could be therefore excluded (Supplementary Figure S11 and S12).

### 3. Electron microscopy and FISH analysis of formed cell aggregates

After about 25 transfers on lactate and PCE, all co-cultures formed spherical aggregates up to 2 mm in diameter (Figure 3A). Field emission-scanning electron microscopy (FE-SEM) was applied to uncover the cell morphology and cell distribution in these aggregates. After preparation for FE-SEM, the sizes of the aggregates were lower (30 to 200 μm, Figure 3B and C). Electron micrographs of both co-cultures revealed a compact network of *S. multivorans* and *D. mccartyi* cells coiled around net-forming filament-like structures. The cells were embedded in an extracellular matrix (ECM) which might aid cell-to-cell contact (Figure 3D, Supplementary Figure S13). *S. multivorans* could be distinguished from *D. mccartyi* by FISH with specific oligonucleotide probes targeting 16S rRNA and 3-dimensional imaging showed a spatial organization and an almost equal distribution of both species within the aggregates (Figure 3E, Supplementary Figure S14). The high resolution of the confocal laser scanning microscopy enabled visualization of single cells and revealed the same morphologies as in the electron micrographs. In the FISH pictures, the sizes of the aggregates ranged from 50 to 100 μm in diameter.

**Figure 3:**
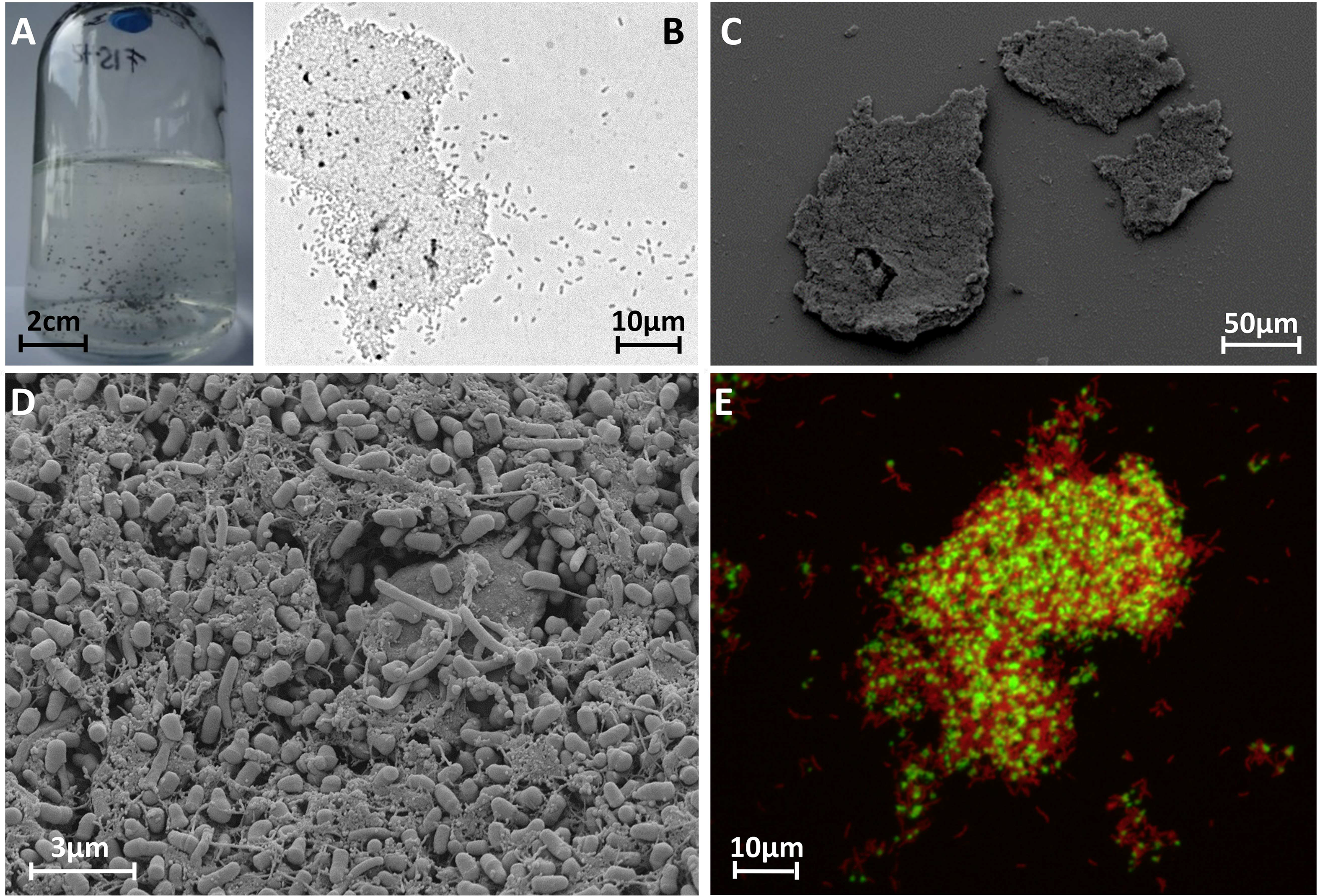
Microscopic analysis of cell aggregates in co-cultures of *S. multivorans* and *D. mccartyi* strains 195 and BTF08. (A) Serum bottle (200 mL) of a *Sm*/BTF08 co-culture showing cell aggregates. (B) Light microscopic image of a *Sm*/BTF08 aggregate. (C, D) Scanning electron micrographs of an aggregate of *Sm*/195. (E) Confocal laser scanning image of FISH stained aggregates of *Sm*/BTF08. Red – *S. multivorans*, green – *D. mccartyi*.

*S. multivorans* revealed a typical helical rod-shaped cell structure and a size of 2 to 5 μm by 0.5 μm at magnifications of around 10.000x and 20.000x in FE-SEM as described previously [55] (Figure 4, Supplementary Figure S13). However, the typical polar flagellum was only observed in a few cells and several flagella seemed to be detached from cells, possibly a part of the ECM (Figure 4, Supplementary Figure S13). *D. mccartyi* showed an atypical cell morphology in the co-cultures. Microscopic analysis of the pure culture revealed a disc-shaped irregular coccus of 0.5 μm diameter (Figure 4a, Supplementary Figure S15), as previously described [11], whereas the *D. mccartyi* strains in the co-culture showed 0.5 μm large barrel-like cells with a flattened cell pole at one side and a ring-shaped septum (Figure 4, Supplementary Figure S13). Visual analysis of cells separated from aggregates in the co-culture identified unequivocally both organisms and confirmed the presence and unusual morphology of *D. mccartyi* (Supplementary Figure S15 and S16).

**Figure 4:**
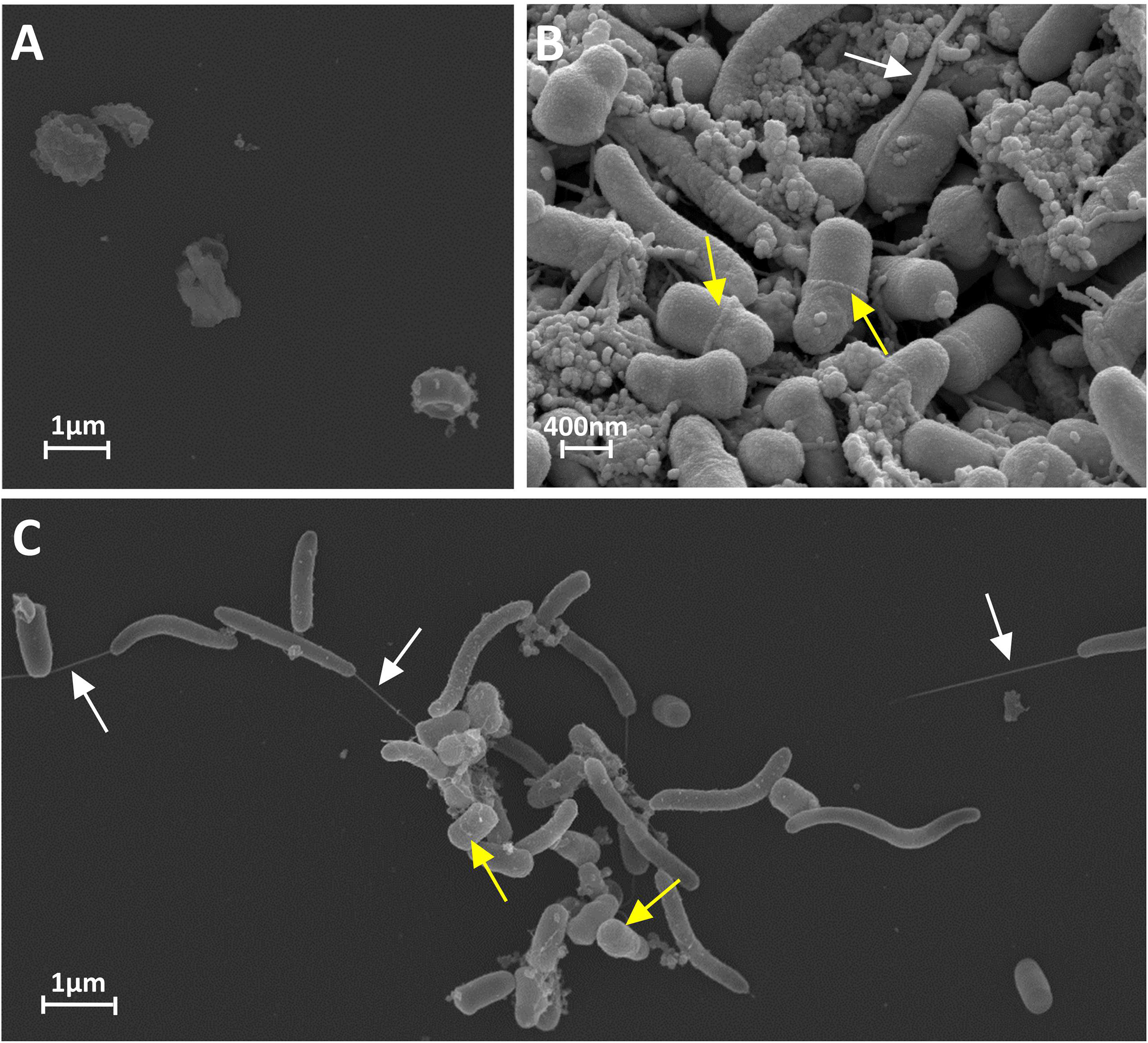
Different cell morphologies of *D. mccartyi* strain BTF08 cells in pure culture (A) and the hitherto unknown barrel-like morphology in co-culture with *S. multivorans* (B, C). White arrows indicate flagella and yellow arrows indicate ring-shaped septum.

### 4. Proteomics of pure and co-cultures

We applied a label-free shotgun proteomics approach to identify changes in protein abundances during PCE dechlorination in pure and co-cultures (with and without vitamin B_12_) with *D. mccartyi* strain BTF08 and *S. multivorans*. The clustering approach nMDS based on quantified proteins revealed a significant separation between pure and co-cultures (*Dhc* BTF08 proteins p=0.001, *Sm* proteins p=0.004, Supplementary Figure S17). A multi-level pattern analysis was applied to determine indicator proteins for a given condition (i.e. co-culture vs. pure culture) to better understand the functional changes occurring during the dehalogenation in the co-culture (Supplementary Excel File S2).

Of the 20 reductive dehalogenases encoded in the genome of *D. mccartyi* strain BTF08 [57], only two RDase proteins were identified, the gene products of btf_1393 (a hitherto not annotated RdhA) and btf_1407 (the vinyl chloride reductase VcrA). The two reductive dehalogenases PceA (PCE reductive dehalogenase) and TceA (TCE reductive dehalogenase), putatively involved in dechlorination of PCE to *c*DCE, were not identified under any condition. While VcrA was one of the most abundant proteins under all tested conditions (Figure 5, Supplementary Figure S18, Supplementary Excel File S1), the gene product of btf_1393 was more abundant in the two co-cultures (Figure 5, Supplementary Excel File S2). A BLASTp search against the NCBI nr database revealed that the btf_1393 amino acid sequence was almost identical (99% or 497/498 amino acid sequence identity over the whole length) to an RdhA from *D. mccartyi* 11a5, encoded by 11a5_1355 and characterized as a novel PCE reductive dehalogenase PteA [56,58]. In *S. multivorans*, the proteins encoded in the organohalide respiratory gene were present among all conditions as described previously [59, 60] (Supplementary Excel File S1).

**Figure 5:**
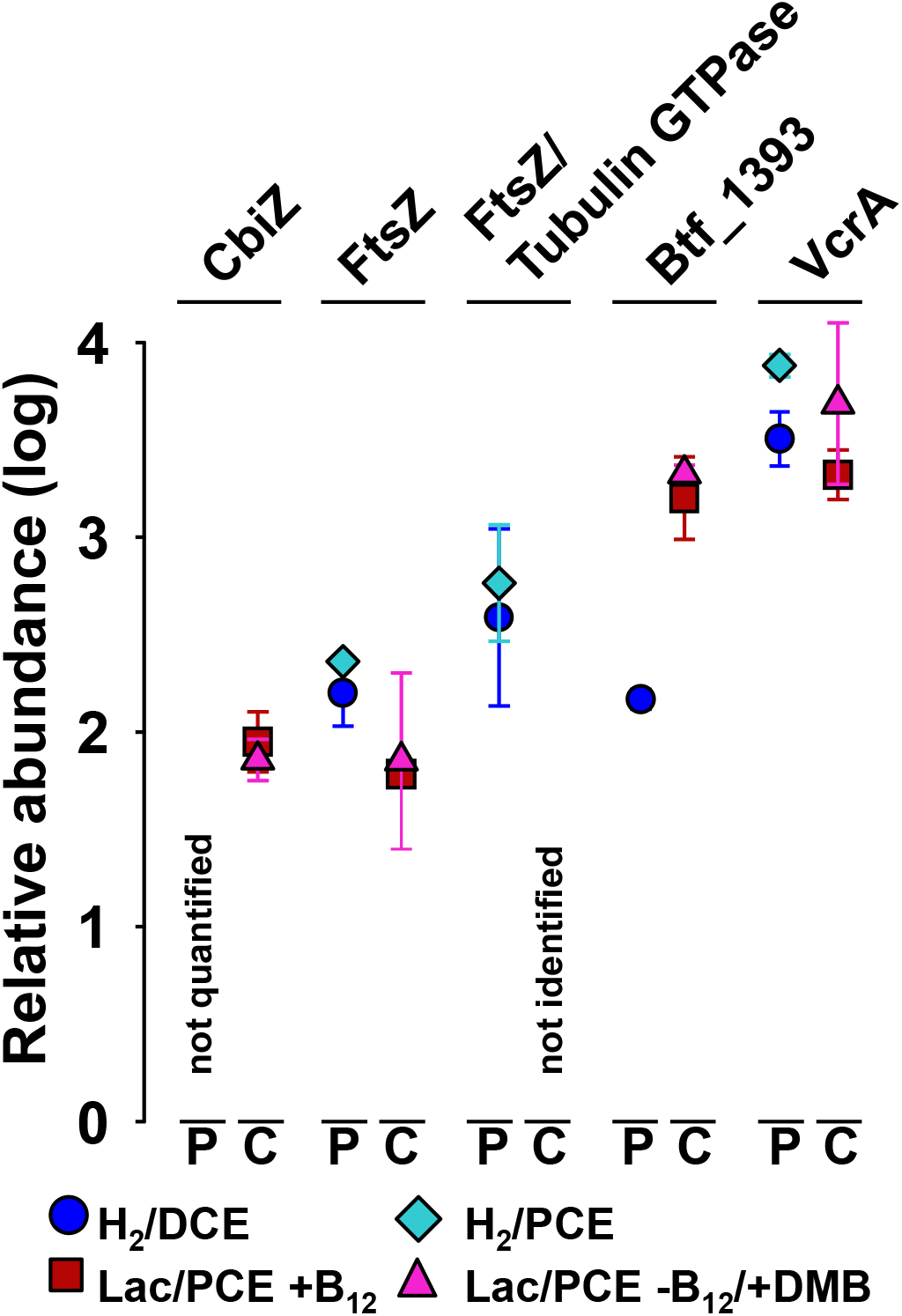
Protein abundances of reductive dehalogenases and a set of significant indicator proteins of *Dhc* BTF08 in pure and co-cultures. Average protein abundance values under different cultivation conditions are shown. Abundances represent log10 fold changes of the median, scaled to zero. The median of all proteins is at 2.3 (see Supplementary Figure S18, which displays each replicate in a scatter plot graph). Four replicates (N=4) were used for proteomic analyses; one of the four replicates were discarded from each of the *D. mccartyi* pure cultures (see Methods). Error bars represent standard deviation, which is covered completely by the symbol if <0.1. P – *D. mccartyi* strain BTF08 pure culture, C – *Sm*/BTF08 co-culture, CbiZ – adenosylcobinamide amidohydrolase (btf_610), FtsZ – cell division protein (btf_0595), FtsZ/Tubulin GTPase (btf_0551), VcrA – vinyl chloride reductive dehalogenase (btf_1407), btf_1393 – reductive dehalogenase homolog to 11a5_1355 of *D. mccartyi* 11a5.

Only a few proteins encoded in the genomes of *S. multivorans* or *D. mccartyi* are annotated to play a potential role in the formation of aggregates, e.g. via an ECM. The proteins which are part of a putative type II pili system in *D. mccartyi* (encoded by btf_1229 to btf_1240) were present in low to medium amounts among all conditions (about median [Med] to lower than Med minus one standard deviation [SD]) but not significantly more abundant in the co-cultures. Many proteins involved in flagellar motility of *S. multivorans* were lower abundant in the co-culture, with most of the Flg and Fli proteins not detected (Supplementary Excel File S1). Protein indicator analysis revealed further that several outer membrane porins of *S. multivorans* (encoded by SMUL_0494, SMUL0693, SMUL_0926 and SMUL_2351) were relevant proteins for the co-cultures. In addition, several molybdopterin oxidoreductases (e.g. a nitrate reductase) of *S. multivorans* were highly abundant only in the co-culture (Supplementary Table S3). Proteins related to cell division (tubulin/GTpase encoded by btf_0551, FtsZ, btf_0595, FtsH, btf_357, a cell division trigger factor, btf_0631) were more abundant or exclusively quantified in pure cultures (Figure 5, Supplementary Table S4, Supplementary Excel File S2). A hypothetical protein encoded close to the tubulin (btf_0548) was also highly abundant (>Med+SD or >Med) only in cells of the pure culture, while it could not be quantified in any of the co-cultures.

The amidohydrolase CbiZ (btf_610), responsible for cleavage of the nucleotide loop of corrinoids, was quantified only in both co-culture conditions in average abundance (Figure 5). The L-threonine 3-O-phosphate decarboxylase CobD, which synthesizes the linker of the lower base and the corrinoid ring of cobamides, could not be quantified in any culture.

## Discussion

In this study, we investigated the dechlorination of PCE to ethene (or vinyl chloride) in co-cultures of *S. multivorans* with *D. mccartyi* BTF08 and 195. Dechlorination profiles, metabolite analysis and growth studies point to a biphasic physiology of the co-culture. In the first stage, when PCE or TCE serve as electron acceptors, *S. multivorans* grows by organohalide respiration with lactate as electron donor. In this phase, *D. mccartyi* does not contribute significantly to PCE dechlorination as indicated by the stable isotope fractionation pattern and the corresponding enrichment factors. This can be explained by the insufficient supply of *D. mccartyi* with hydrogen as electron donor due to the lack of (or very low) hydrogen production by *S. multivorans* during respiratory growth. In the second phase of the co-culture, when PCE has been completely converted to *c*DCE after less than two days, *S. multivorans* grows via fermentation of lactate mainly to acetate, CO2 and hydrogen [36]. The fate of a part of the organic carbon and of the electrons generated by lactate oxidation could not be elucidated in this study. This part might have been used for biomass production and the generation of small amounts of organic acids, such as succinate, below the detection limit of the HPLC used here. It has been shown earlier that, for thermodynamic reasons, lactate can only be utilized fermentatively by *S. multivorans* in co-cultures, where hydrogen concentration is kept low by a syntrophic hydrogen consumer, while *S. multivorans* does not grow fermentatively on lactate in pure cultures [36]. In the co-culture described here, *D. mccartyi* utilizes H2 produced by *S. multivorans* as electron donor for reductive dehalogenation of *c*DCE or VC (Figure 6). As expected, PCE was completely dechlorinated to ethene in the *Sm*/BTF08 co-culture and mainly to VC and minor amounts of ethene in *Sm*/195. A similar slow and incomplete VC dechlorination was found for pure cultures of the latter strain [61] and co-cultures of *D. mccartyi* strain 195 with either *Desulfovibrio vulgaris* Hildenborough or *Syntrophomonas wolfei* [21, 22]. *c*DCE was presumably converted to ethene by VcrA of BTF08. This is supported by the proteomic analysis described here, in which VcrA of *D. mccartyi* strain BTF08 was highly abundant and the predominant RDase in all cultures. A highly similar VcrA ortholog of *D. mccartyi* strain VS (99% amino acid sequence identity) has already been biochemically characterized and described to dechlorinate *c*DCE and VC [62]. Interestingly, neither TceA, nor PceA, which had been suggested to be responsible for dechlorination of PCE to VC [57], were detected in any of the samples. An identical ortholog of the only other detected RDase (encoded by btf_1393) was recently characterized as PteA, a novel PCE reductive dehalogenase dechlorinating PCE to TCE in *D. mccartyi* strain 11a5 [58]. *D. mccartyi* strain 195 was not subject to proteomic investigation in our study. However, previous studies suggested that TceA of this organism is involved in *c*DCE to VC dechlorination and that VC is further dechlorinated slowly by the same enzyme [61, 63].

**Figure 6:**
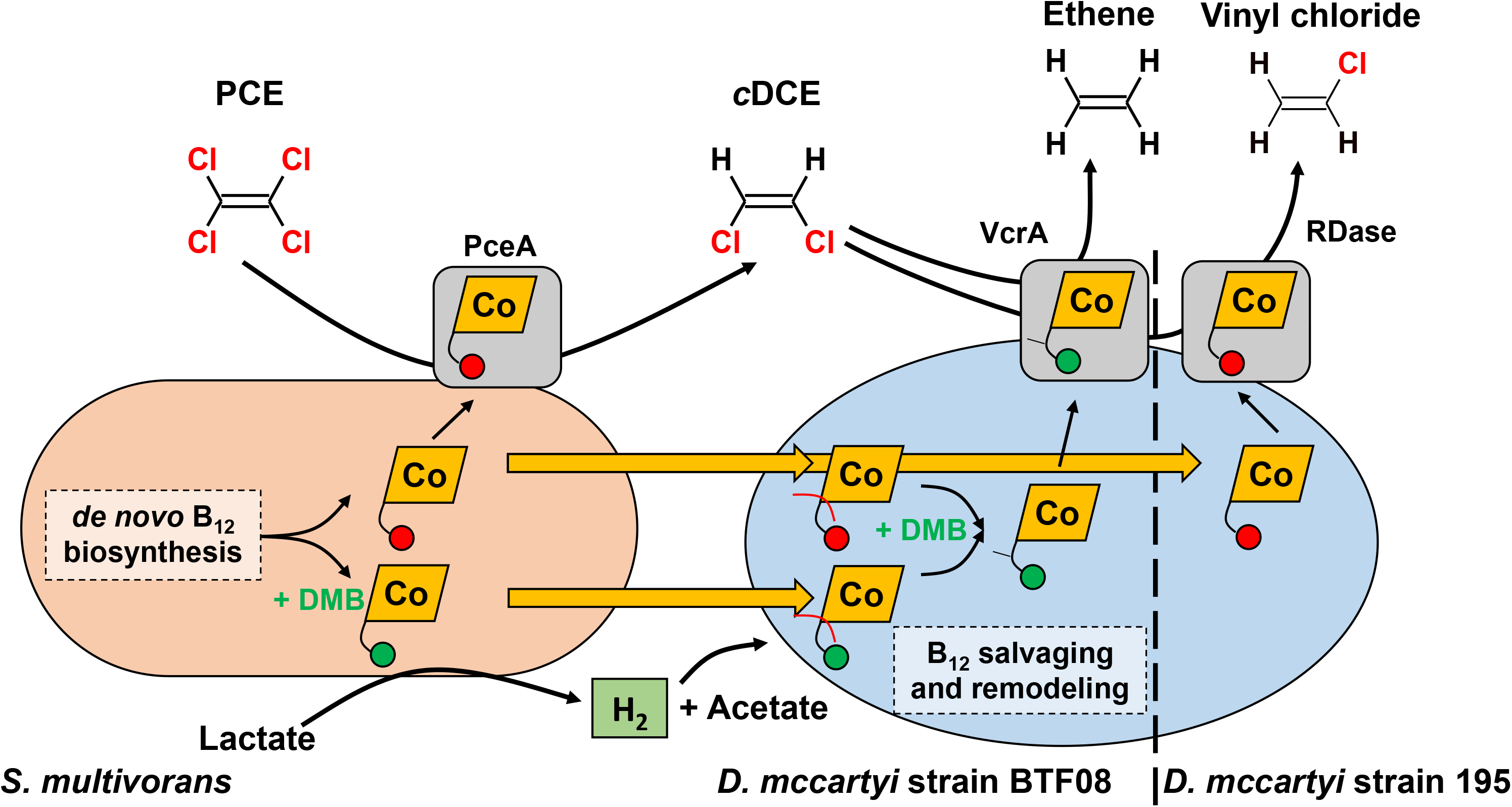
Interspecies metabolite transfer of *Sulfurospirillum multivorans* and *Dehalococcoides mccartyi*. PCE is dechlorinated to *c*DCE by *S. multivorans* with electrons from lactate oxidation. After depletion of PCE, *S. multivorans* switches to fermentative metabolism, thereby generating hydrogen, which is consumed by *D. mccartyi* as electron donor. The electron acceptor for *D. mccartyi* is *c*DCE and is further dechlorinated to ethene or VC by *D. mccartyi*. *S. multivorans* synthesizes norpseudo B_12_ ([Ade]NCba, 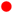) and [DMB]NCba 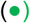 when DMB (maximally 1 μM) is amended. Both can be salvaged and remodeled by *D. mccartyi* into [DMB]Cba (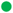 with a y-shaped linker). The interspecies cobamide transfer is indicated by yellow arrows. A discrimination between norcobamide or cobamide incorporation into the *D. mccartyi* RDases is not depicted here, since the data do not allow a conclusion on the cobalamin nucleotide loop type.

### Norpseudo-B_12_, produced by *S. multivorans*, is used differently by the two *D. mccartyi* strains

Without amendment of vitamin B_12_ the *Sm*/BTF08 co-culture was not capable of *c*DCE dechlorination to ethene, indicating that *D. mccartyi* strain BTF08 cannot use the norpseudo-B_12_ ([Ade]NCba) synthesized by *S. multivorans* for *c*DCE dechlorination. Complete dechlorination to ethene by *D. mccartyi* strain BTF08 was restored by the addition of the lower ligand DMB, indicating that the non-functional adenine ligand of [Ade]NCba could be replaced by DMB. This assumption is supported by a previous study, in which *D. mccartyi* was shown to incorporate different lower ligands into cobamides [17]. Therefore, it was not surprising that DMB-containing cobalamin were detected in the *Sm/Dhc* co-cultures. Most probably, after addition of DMB the detected [DMB]NCba was synthesized by *S. multivorans* besides [Ade]NCba, since the organism is able to incorporate different benzimidazoles and generate the corresponding cobamides [35, 64]. [DMB]Cba could then be produced during salvaging and remodeling of available [Ade]NCba and/or [DMB]NCba by *D. mccartyi*. It can, however, not be excluded from our data that [DMB]NCba can be used as well by *D. mccartyi* strain BTF08 for cDCE or VC dechlorination. The nucleotide loop cleavage is most likely mediated by the adenosylcobinamide hydrolase CbiZ, as shown for CbiZ of *Rhodobacter sphaeroides* [65, 66]. Of the two CbiZ encoded in the genome of *D. mccartyi* strain BTF08, one was detected in the proteome and showed a higher abundance in the co-culture compared to the pure culture. These results present evidence of the involvement of CbiZ in exchanging the complete nucleotide loop in *D. mccartyi*.

Different from the *c*DCE accumulation in *Sm*/BTF08 co-cultures grown without DMB, the *Sm*/195 coculture dechlorinated PCE to VC in media without vitamin B_12_ or DMB. This indicates *c*DCE to VC conversion by strain 195 with [Ade]NCba produced by *S. multivorans*. However, previous studies showed [Ade]Cba (or other cobalamins with adenine as lower ligand) to be non-functional in *D. mccartyi* strain 195 with TCE as electron acceptor [17]. Trace amounts of DMB or cobalamin could be an explanation, but this explanation is unlikely given the same treatment but different dechlorination pattern of the co-cultures. Since RDase gene expression is often dependent on the electron acceptor present [14], *c*DCE in the *Sm*/195 co-culture might have induced a yet unknown RDase dechlorinating *c*DCE to VC with [Ade]NCba as cofactor. Alternatively, TceA containing [Ade]NCba could catalyze cDCE to VC dechlorination, but not TCE dechlorination to *c*DCE. The mechanism of cobalamin transfer from *S. multivorans* to *D. mccartyi* is not yet known. Most likely, cobalamin is released upon cell lysis or degradation of periplasmic PceA and possibly excreted through porins of *S. multivorans*, some of which were higher abundant in co-cultures.

### Cell aggregates formed by *D. mccartyi* and *S. multivorans* could enhance interspecies hydrogen transfer

The cell aggregates observed in all studied co-cultures but not in pure cultures are common for obligate syntrophic interactions often found in e.g. acetogenic and methanogenic communities [67, 68, 69]. The decrease of intermicrobial distances and establishment of cell-to-cell contacts, which were observed in co-cultures via FE-SEM, should lead to increased metabolite (e.g. hydrogen) fluxes between species according to Fick’s law [70], ultimately enhancing growth and dechlorination rates. Experimentally, this was shown, for example, during syntrophic propionate conversion of *Pelotomaculum thermopropionicum* SI and *Methanothermobacter thermoautotrophicus* ∆H where interspecies hydrogen transfer was calculated to be optimal in aggregates [71]. EPS-like substances and flagella most likely contribute to a stabilization of aggregates by adhesion and attachment of the cells [67, 72]. In addition, cellular extrusions such as pili and flagella might contribute to direct interspecies electron transfer (DIET). While *S. multivorans* produces flagella in pure cultures [55], several structural flagellar proteins were down-regulated in the co-culture, speaking against a use of flagellar for DIET in the co-culture of *D. mccartyi* and *S. multivorans*. Other proteins putatively involved in DIET are not encoded in the genomes of *D. mccartyi* or *S. multivorans*, arguing for an exclusive interspecies transfer of electrons via hydrogen. Aggregated *D. mccartyi* cells embedded in an ECM were observed in a bioreactor community dechlorinating trichloroethene to ethene [73]. However in this study, *D. mccartyi* formed typical disc-shaped cells, in contrast to the unusual, never before observed barrel-like morphology of *D. mccartyi* in the co-cultures investigated in our study. This barrel-like morphology might be caused by the down-regulation of proteins involved in cell division in the co-culture. One of these proteins, FtsZ, localized at the cell division site, was shown to play a key role in cytokinesis in *E. coli*. It was shown to be responsible for septal invagination of the cell wall and cytoplasmic membrane by forming a ring-shaped septum followed by cell division [74, 75, 76]. Its down-regulation might hamper a complete membrane constriction resulting in slower cell division that might cause the observed barrel-like cell morphologies. However, the reasons for this down-regulation and the different morphology remain obscure. Possibly the less frequent cell divisions and/or the barrel-like morphology give *D. mccartyi* an advantage in cell-to-cell-contact. In addition, this shape could represent a “sessile” state in which *D. mccartyi* adjusts its metabolism to the given condition in the aggregates, including a downregulation of cell division. Since electron micrographs of co-cultures with *D. mccartyi* are scarce, it is not possible to state whether the unusual cell morphology is specific for the partnership with *S. multivorans* or whether it is found frequently for co-cultures containing *D. mccartyi*. With *S. wolfei* as syntrophic partner, disc-like cell structures typical for *D. mccartyi* strain 195 were observed [21].

## Conclusion

This study provides first insights into the interactions of *S. multivorans* in association with *D. mccartyi* and the formation of aggregates in which *D. mccartyi* showed an unusual barrel-like morphology. Dechlorinating microbial communities sometimes reveal the presence of *Sulfurospirillum* spp., but the functional role of these Campylobacterota (formerly Epsilonproteobacteria) is unexplored [77]. We observed that interspecies hydrogen and cobamide transfer in the co-culture resulted in fast and complete dechlorination of PCE to ethene. *S. multivorans* could provide all growth factors required by *D. mccartyi*, including hydrogen and *c*DCE as energy source, acetate as carbon source and cobamides as RDase cofactors (Figure 6). It is the first study in which *D. mccartyi* was provided with all nutrients required for growth by its syntrophic partner. Additionally, PCE to *c*DCE dechlorination was sped up by the fast dechlorination rate of *S. multivorans*. This is of high interest for bioremediation attempts using *Dehalococcoides-containing* mixed cultures, since electron donor and cobalamin limitations may impede *Dehalococcoides* dechlorination activities.

## Acknowledgement

This work was supported by the Jena School for Microbial Communication (JSMC) and the DFG Research Unit FOR 1530 (sub-projects 1, 2, 5 and 6 were involved). The authors would like to thank Benjamin Scheer and Kathleen Eismann (Helmholtz Centre for Environmental Research, Leipzig) for skillfull technical assistance in the lab, Susanne Linde (University Hospital Jena, Center for Electron Microscopy) for the field-emission scanning electron microscopic analysis and Dr. Elena Romano (University of Rome Tor Vergata, Department of Biology) for invaluable assistance in the confocal laser scanning microscope image acquisition and processing. Dr. Torsten Schubert (Friedrich Schiller University Jena) is gratefully acknowledged for insightful discussions on corrinoid metabolism.

