## Supplementary Information for "Interspecies metabolite transfer and aggregate formation in a co-culture of *Dehalococcoides* and *Sulfurospirillum* dehalogenating tetrachloroethene to ethene"

**Stefan Kruse[1], Dominique Türkowsky[2], Jan Birkigt[3], Bruna Matturro[4], Steffi Franke[3], Nico Jehmlich[2], Martin von Bergen[2,6], Martin Westermann[5], Simona Rossetti[4], Ivonne Nijenhuis[3], Lorenz Adrian[3,7], Gabriele Diekert[1], Tobias Goris[1,8]**

[1]Department of Applied and Ecological Microbiology, Institute of Microbiology, Friedrich Schiller University, Jena, Germany

[2]Department Molecular Systems Biology, Helmholtz Centre for Environmental Research – UFZ, Leipzig, Germany

[3]Department of Isotope Biogeochemistry, Helmholtz Centre for Environmental Research – UFZ, Leipzig, Germany

[4]Water Research Institute, IRSA-CNR, Monterotondo, Rome, Italy

[5]Center for Electron Microscopy of the University Hospital Jena, Jena, Germany

[6]Institute of Biochemistry, Faculty of Life Sciences, University of Leipzig, Leipzig, Germany

[7]Chair of Geobiotechnology, Technische Universität Berlin, Germany

[8]Current address: Department of Molecular Toxicology, Research Group Intestinal Microbiology, German Institute of Human Nutrition Potsdam-Rehbruecke, Nuthetal, Germany

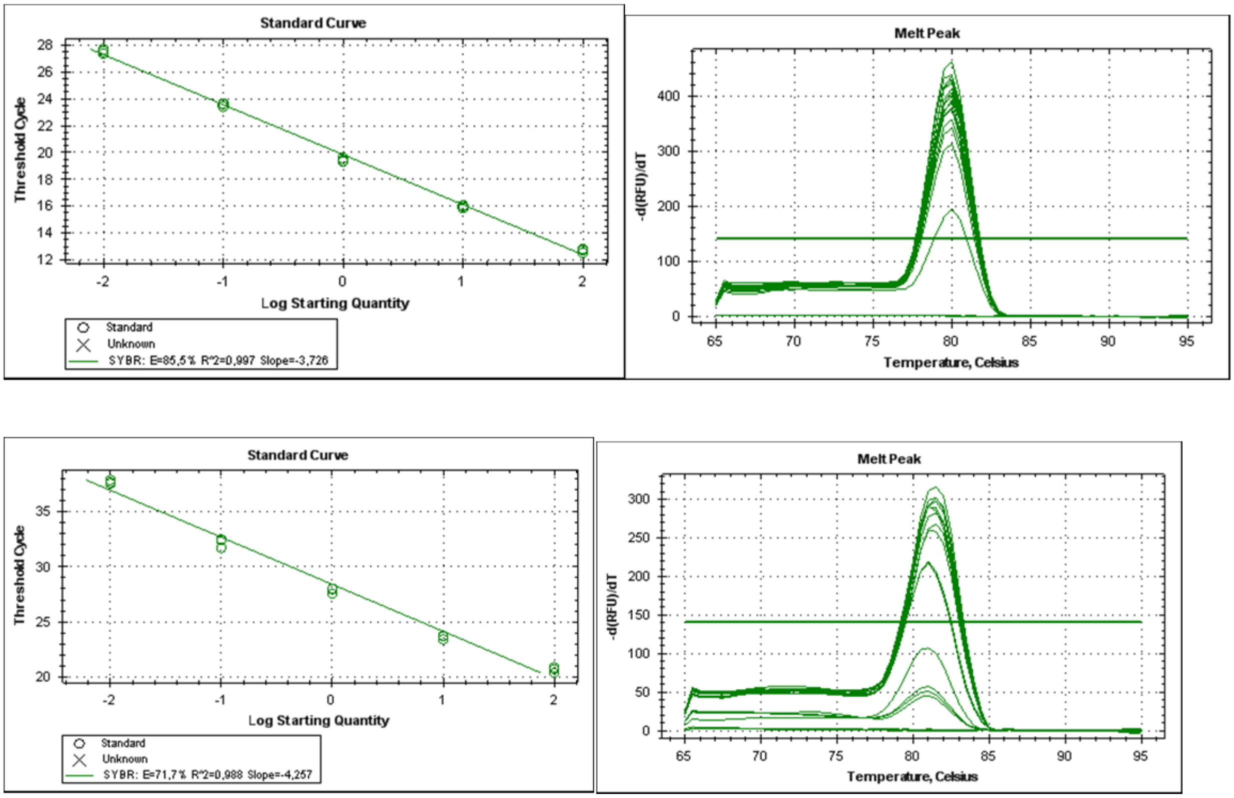


**Supplementary Figure S1: Melting standard curves and primer melting curves of qPCR.** *S. multivorans* figures on top (Log Starting Quantity of 0 = 1x$10^7$ cells), *D. mccartyi* BTF08 (Log Starting Quantity of 0 = 1x$10^6$ cells) on the bottom.

**Supplementary Table S1: Calculation of theoretical masses of the isotope distribution of different corrinoid types.** Values according to IDCalc.

| $B_{12}$ type | Lower ligand | Linker | Formula | Rel. abundance | m/z z = 2 |
|---|---|---|---|---|---|
| **Cyanocobalamin [DMB]Cba** | DMB | Aminopropan-2-ol *O*-2-phosphate | $C_{63}H_{88}N_{14}O_{14}PCo$ | 100.00<br>75.98<br>31.35<br>9.20<br>2.13 | 678.2910<br>678.7922<br>679.2933<br>679.7945<br>680.2957 |
| **5-methoxy-benzimidazolyl-cobamide [5-OMeBza]Cba** | 5-OMeBza | Aminopropan-2-ol *O*-2-phosphate | $C_{61}H_{84}N_{14}O_{15}PCo$ | 100.00<br>73.77<br>29.91<br>8.68<br>2.00 | 672.2728<br>672.7740<br>673.2751<br>673.7763<br>674.2775 |
| **Norpseudo-vitamin $B_{12}$ [Ade]NCba** | Adenine | Ethanolamine *O*-phosphate | $C_{58}H_{81}N_{17}O_{14}PCo$ | 100.00<br>71.51<br>28.07<br>7.89<br>1.76 | 665.7682<br>666.2694<br>666.7706<br>667.2717<br>667.7729 |
| **Norvitamin $B_{12}$ [DMB]NCba** | DMB | Ethanolamine *O*-phosphate | $C_{62}H_{86}N_{14}O_{14}PCo$ | 100.00<br>74.85<br>30.50<br>8.85<br>2.03 | 671.2832<br>671.7843<br>672.2855<br>672.7867<br>673.2879 |

**Supplementary Table S2: Comparison of theoretical masses of different vitamin $B_{12}$ types and detected masses in the co-cultures of *Sm*/195, *Sm*/BTF08 and pure cultures of *Dhc* strain 195.** Theoretical masses were obtained from Supplementary Table S1.

| **Theoretical masses of vitamin $B_{12}$ types** | | | **MS detected masses [m/z; z = 2]** Samples | | | | | |
|---|---|---|---|---|---|---|---|---|
| **$B_{12}$ type** | Rel. abundance | m/z z = 2 | Vitamin $B_{12}$ standard[1] | *Sm*/195 -$B_{12}$/+DMB | *Sm*/BTF08 -$B_{12}$/+DMB | *Sm*/195 -$B_{12}$/-DMB | *Dhc* 195 +[5-OMe Bza]Cba | *Dhc* 195 +[Ade]NCba |
| **Cyanocobalamin [DMB]Cba** | 100.00<br>75.98<br>31.35<br>9.20<br>2.13 | 678.2910<br>678.7922<br>679.2933<br>679.7945<br>680.2957 | 678.2928<br>678.7892<br>679.2866<br>679.7867<br>680.2886 | 678.2927<br>678.7913<br>679.2912<br>679.7927<br>680.2939 | 678.2904<br>678.7919<br>679.2934<br>679.7949<br>- | - | - | - |
| **5-methoxy-benzimidazolyl-cobamide [5-OMeBza]Cba** | 100.00<br>73.77<br>29.91<br>8.68<br>2.00 | 672.2728<br>672.7740<br>673.2751<br>673.7763<br>674.2775 | - | - | - | - | 672.2745<br>672.7737<br>673.2739<br>673.7755<br>674.2765 | - |
| **Norpseudovitamin $B_{12}$ [Ade]NCba** | 100.00<br>71.51<br>28.07<br>7.89<br>1.76 | 665.7682<br>666.2694<br>666.7706<br>667.2717<br>667.7729 | - | 665.7689<br>666.2698<br>666.7712<br>667.2729<br>667.7739 | 665.7679<br>666.2693<br>667.7706<br>667.1761<br>667.7712 | 665.7698<br>666.2686<br>666.7684<br>667.2699<br>667.7721 | - | 665.7704<br>666.2685<br>666.7681<br>667.2699<br>667.7716 |
| **Norvitamin $B_{12}$ [DMB]NCba** | 100.00<br>74.85<br>30.50<br>8.85<br>2.03 | 671.2832<br>671.7843<br>672.2855<br>672.7867<br>673.2879 | - | 671.2848<br>671.7847<br>672.2855<br>672.7879<br>673.28883 | 671.2833<br>671.7842<br>672.2856<br>672.7867<br>673.2877 | - | - | - |

[1]Mass spectrometric analysis of the vitamin $B_{12}$ standard is given in Supplementary Figure S4

### Vitamin $B_{12}$ standard ([DMB]Cba)

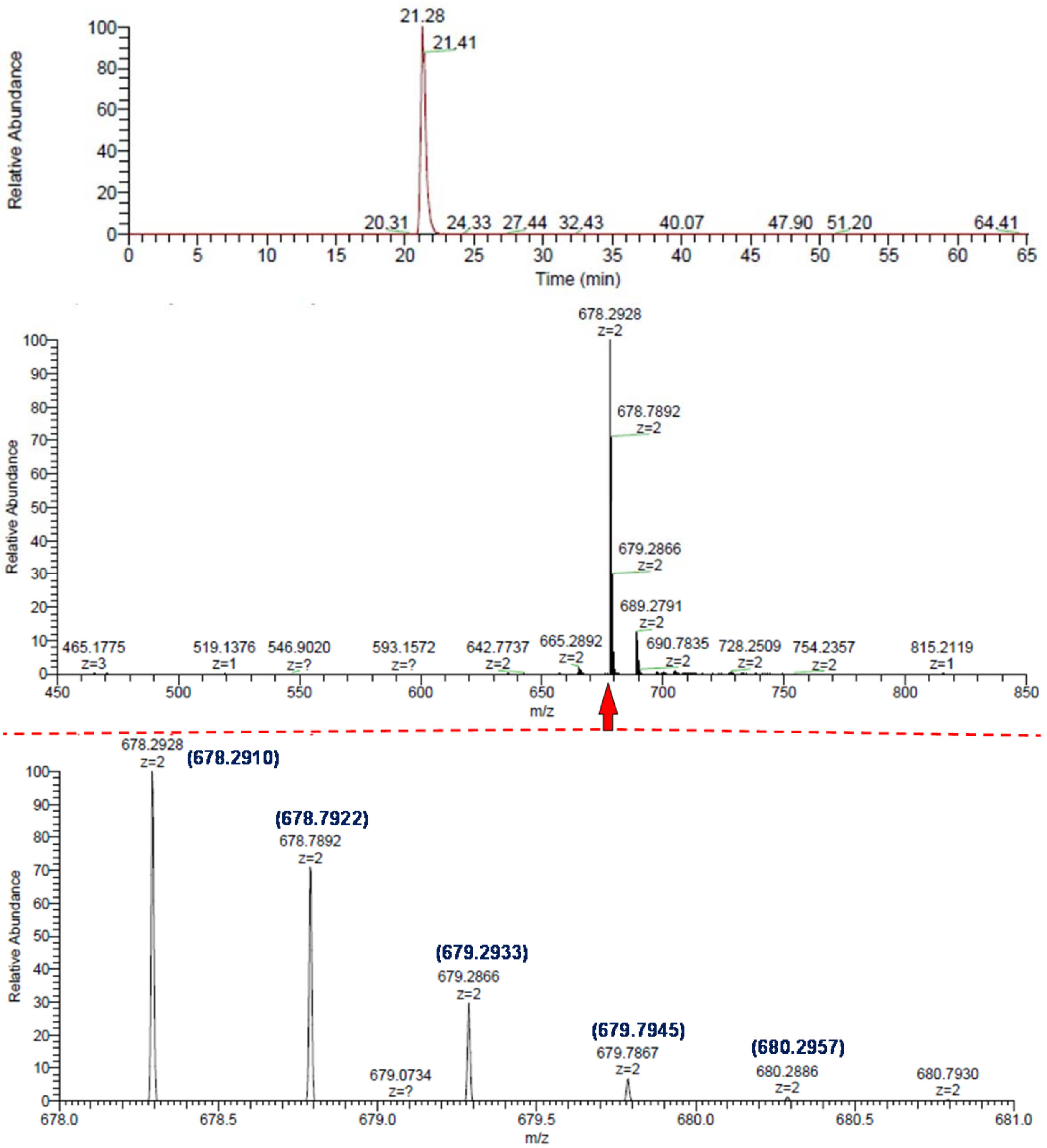


**Supplementary Figure S2: Mass spectrometric analysis of vitamin $B_{12}$ standard ([DMB]Cba, 50 mg/L).** Arrow indicates magnifications of the mass spectrum. Numbers in brackets represent theoretical mass values of corrinoids according to calculated isotope pattern (see Supplementary Table S1 and S2).

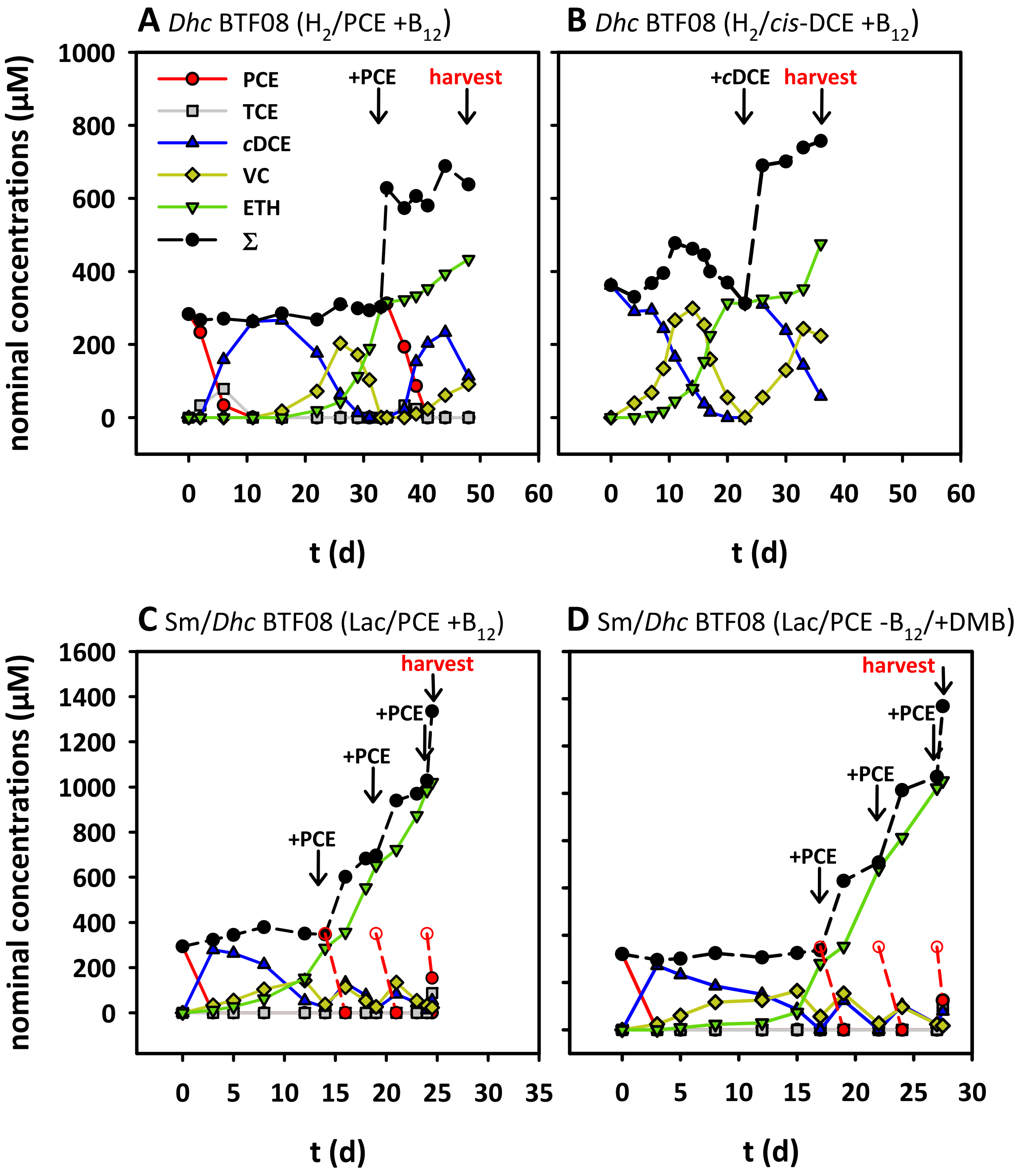


**Supplementary Figure S3: Dechlorination profiles of *Dhc* BTF08 pure culture and *Sm*/BTF08 co-culture used for comparative proteome study.** Depicted graphs are representatives of different biological independent replicates (N(P): 3; N(C): 3; N(L): 4; N(D): 4). Re-fed PCE is indicated by the theoretical amount added (dashed, red line).

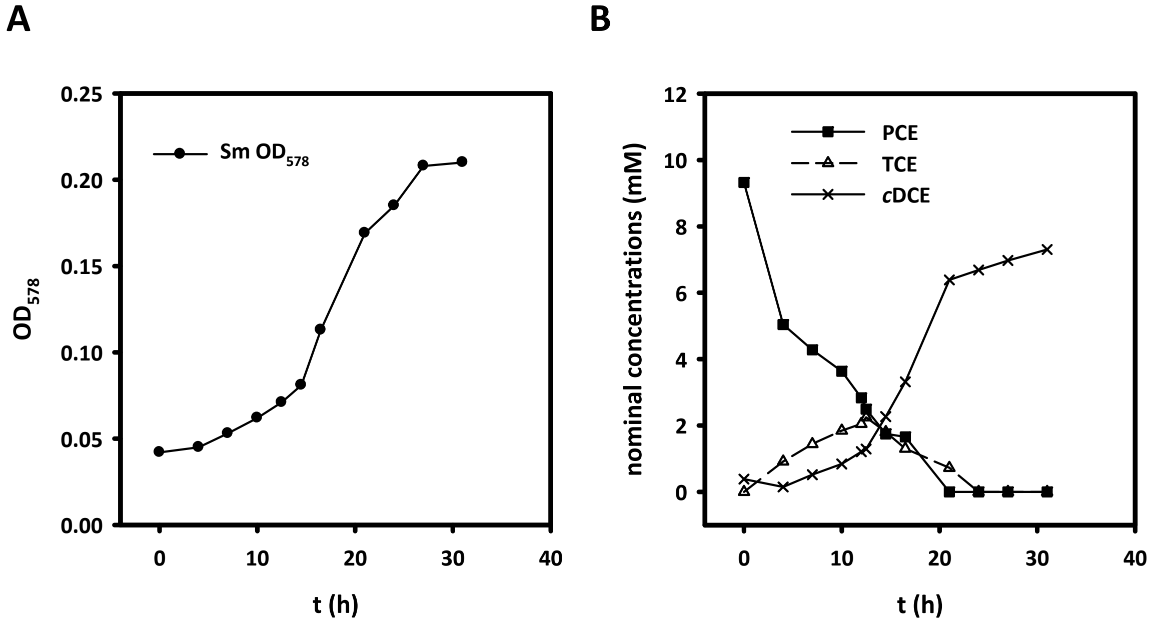


**Supplementary Figure S4: Dechlorination of PCE by *S. multivorans* cultivated in the co-culture medium.** (A) Growth curve. (B) Dechlorination profile. PCE - tetrachloroethene, TCE - trichloroethene, cDCE - cis-1,2-dichloroethene. Lactate (40 mM) was used as electron donor.

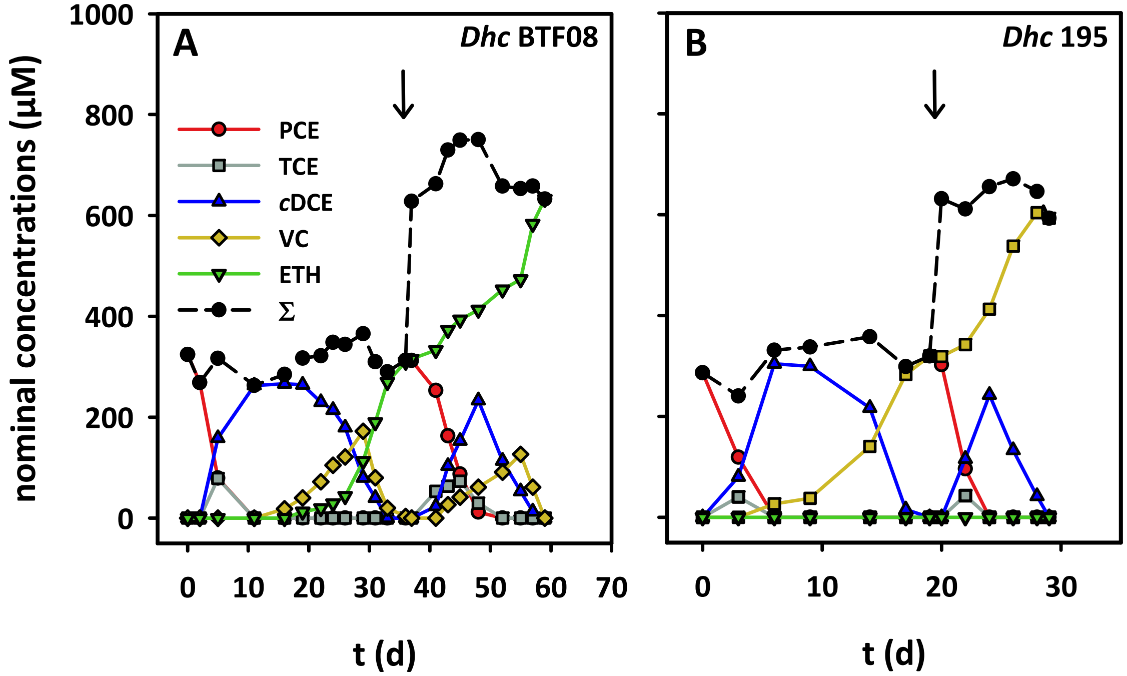


**Supplementary Figure S5: Dechlorination of PCE with $H_2$ as electron donor by pure cultures of (A) *D. mccartyi* strain BTF08 and (B) *D. mccartyi* strain 195.** Arrow indicates refeeding of PCE. ∑ = mass balance; sum of PCE, TCE, *cis*-DCE, VC and ethene.

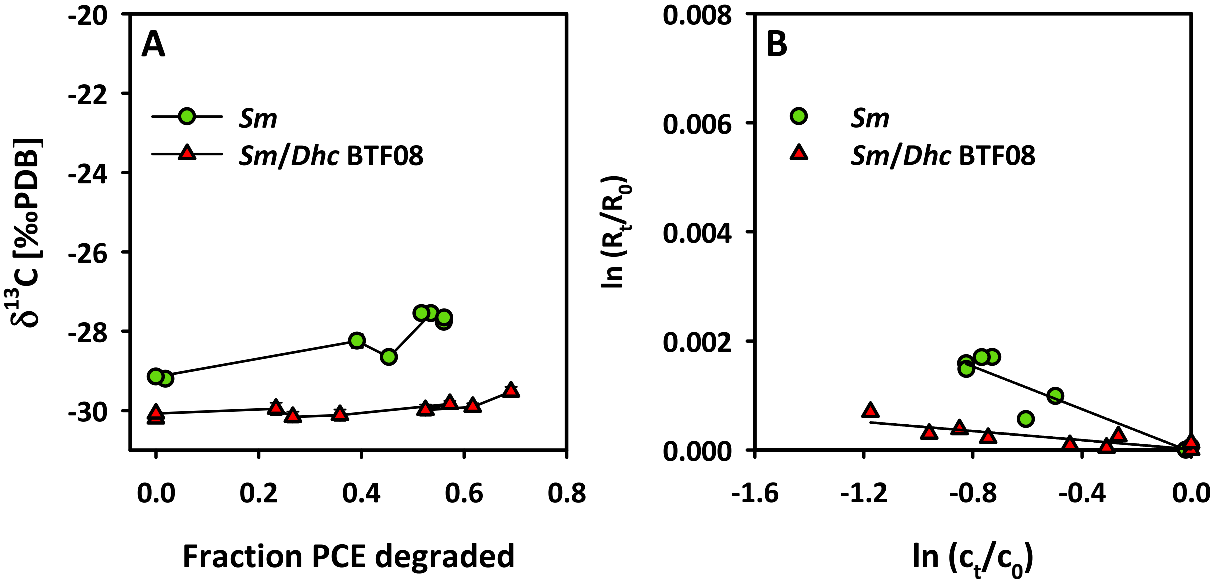


**Supplementary Figure S6: Change in carbon isotope composition (A) and stable isotope fractionation (B) during reductive dechlorination of PCE by *S. multivorans* and the *Sm*/BTF08 co-culture.** (A) SD was calculated from three technical replicates and with <0.5 δ-units covered by the symbols. (B) Correlation of stable isotope fractionation is $R^2$ = 0.859 for *S. multivorans* and $R^2$ = 0.684 for *Sm*/BTF08 co-culture.

**Co-culture *S. multivorans*/*D. mccartyi* strain BTF08 without $B_{12}$ and with DMB**

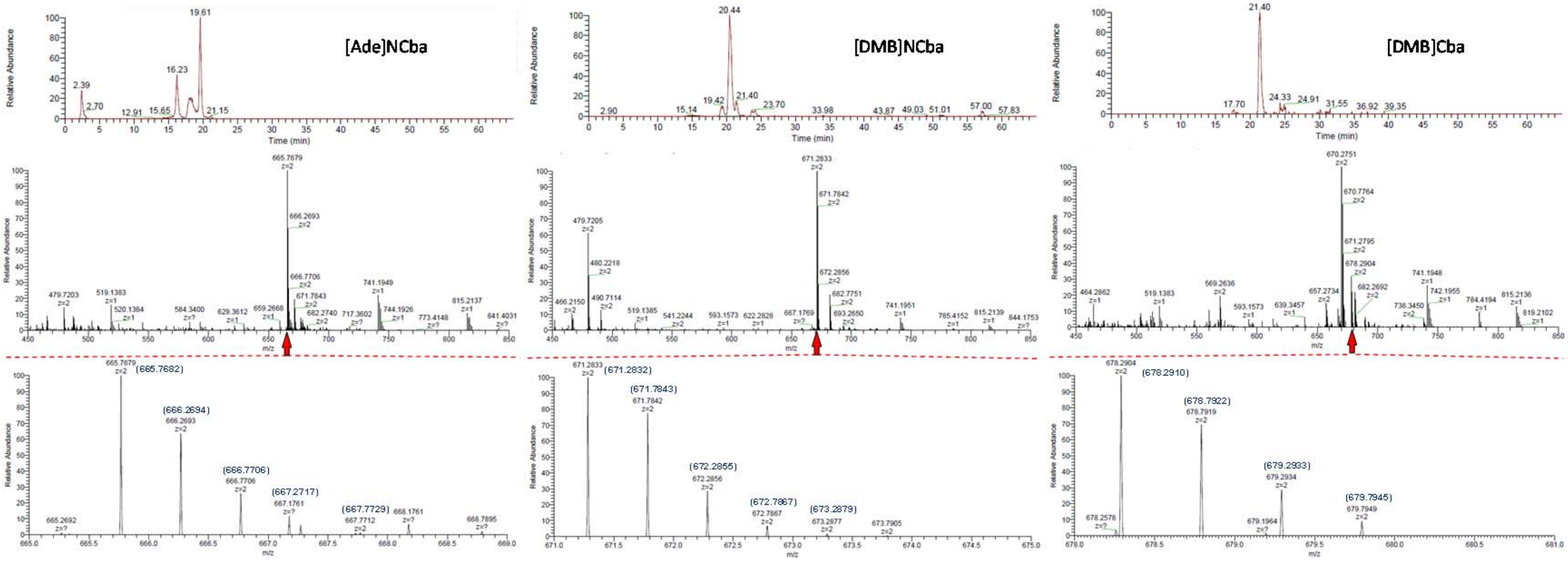


**Supplementary Figure S7: Mass spectrometric analysis of corrinoids extracted from *Sm*/BTF08 co-culture grown on Lac/PCE with 1 µM DMB without amendment of vitamin $B_{12}$.** Arrows indicate magnifications of mass spectra. Numbers in brackets represent theoretical mass values of corrinoids according to calculated isotope pattern (see Supplementary Table S1 and S2). [Ade]NCba - adeninyl-norcobamide, [DMB]NCba - 5,6-dimethylbenzimidazolyl-norcobamide, [DMB]Cba - 5,6-dimethylbenzimidazolyl-cobamide.

### Co-culture *S. multivorans*/*D. mccartyi* strain 195 without $B_{12}$ and without DMB

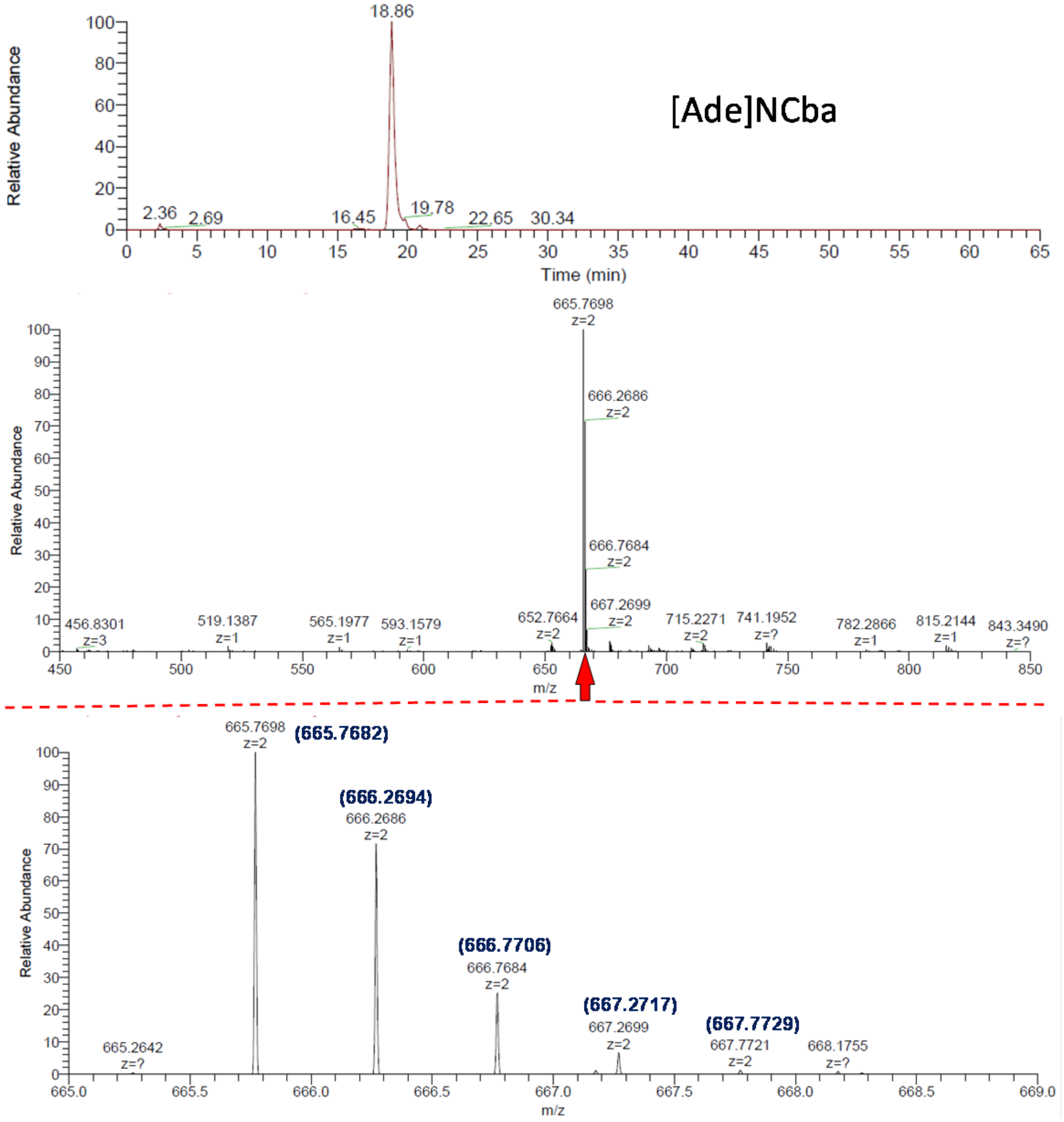


**Supplementary Figure S8: Mass spectrometric analysis of corrinoids extracted from *Sm*/195 co-culture grown on Lac/PCE without amendment of vitamin $B_{12}$and without DMB.** Arrows indicate magnifications of mass spectra. Numbers in brackets represent theoretical mass values of corrinoids according to calculated isotope pattern (see Supplementary Table S1 and S2). [Ade]NCba - adeninyl-norcobamide.

### Co-culture *S. multivorans*/*D. mccartyi* strain 195 without $B_{12}$ and with DMB

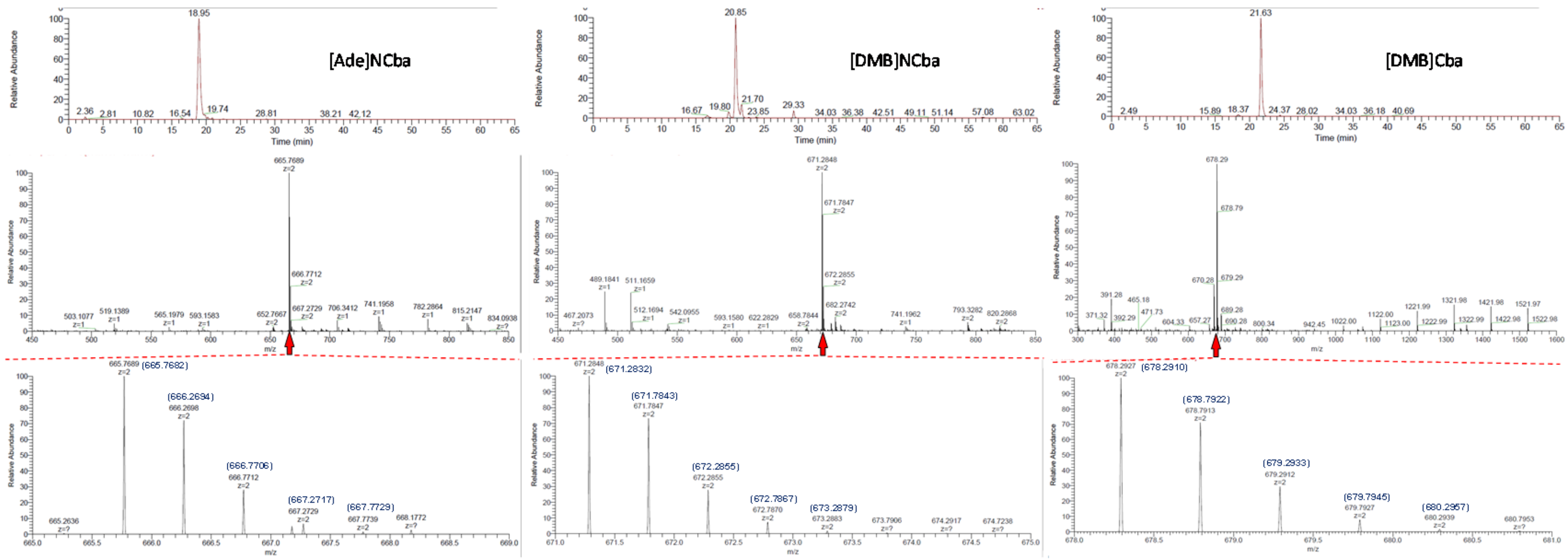


**Supplementary Figure S9: Mass spectrometric analysis of corrinoids extracted from *Sm*/*Dhc* 195 co-culture grown on Lac/PCE with 1 µM DMB without amendment of vitamin $B_{12}$.** Arrows indicate magnifications of mass spectra. Numbers in brackets represent theoretical mass values of corrinoids according to calculated isotope pattern (see Supplementary Table S1 and S2). [Ade]NCba - adeninyl-norcobamide, [DMB]NCba - 5,6-dimethylbenzimidazolyl-norcobamide, [DMB]Cba - 5,6-dimethylbenzimidazolyl-cobamide.

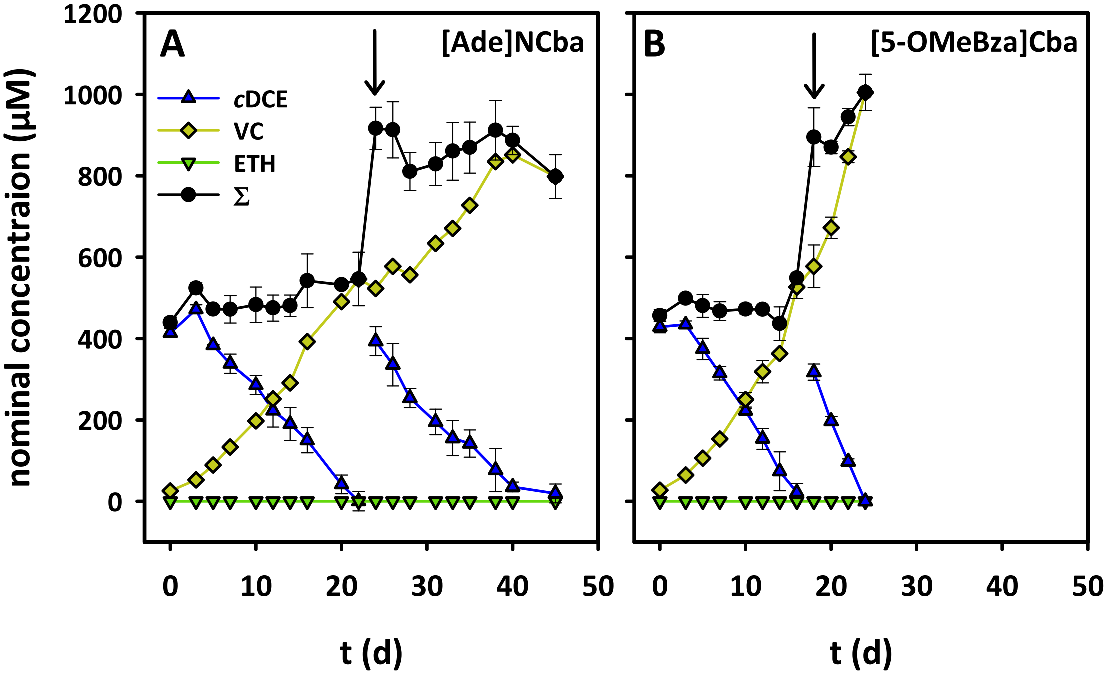


**Supplementary Figure S10: Dechlorination of *cis*-DCE with $H_2$ as electron donor by pure cultures of *D. mccartyi* strain 195 with (A) norpseudovitamin $B_{12}$ and (B)5-methoxybenzimidazolylcobamide .**Arrow indicates refeeding of *cis*-DCE. ∑ = mass balance; sum of *cis*-DCE, VC and ethene. Ade – adenine, NCba – norpseudocobamide, 5-OMeBza - 5-methoxybenzimidazol, , Cba - cobamide.

### Pure culture *D. mccartyi* strain 195 with [5-OMeBza]

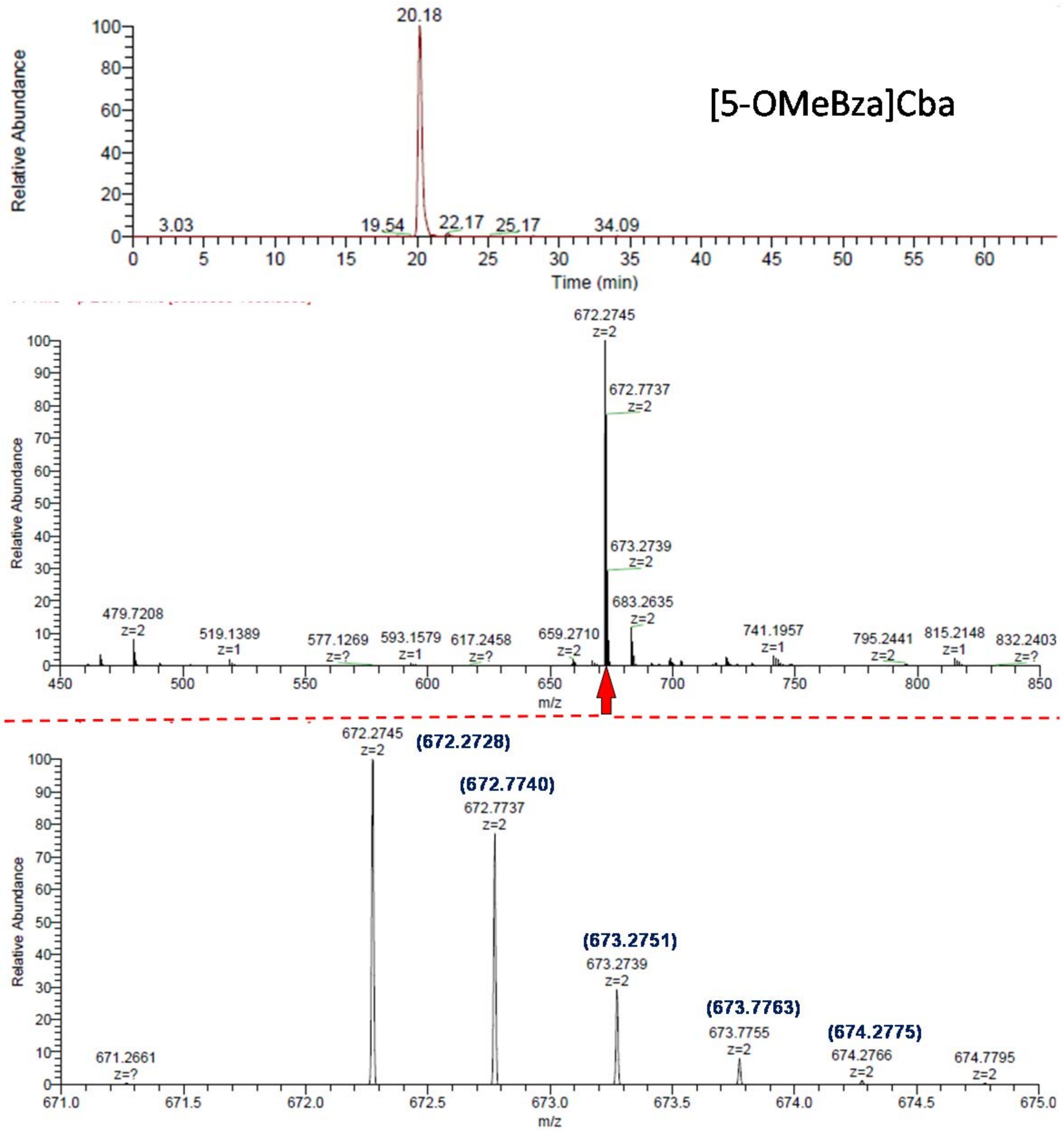


**Supplementary Figure S11: Mass spectrometric analysis of corrinoids extracted from *D. mccartyi* strain 195 grown on $H_2$/*c*DCE with 52 nM [5-OMeBza]Cba.** Arrows indicate magnifications of mass spectra. Numbers in brackets represent theoretical mass values of corrinoids to according calculated isotope pattern (see Supplementary Table S1 and S2). [5-OMeBza]Cba - 5-methoxybenzimidazolyl-cobamide.

### Pure culture *D. mccartyi* strain 195 with [Ade]NCba

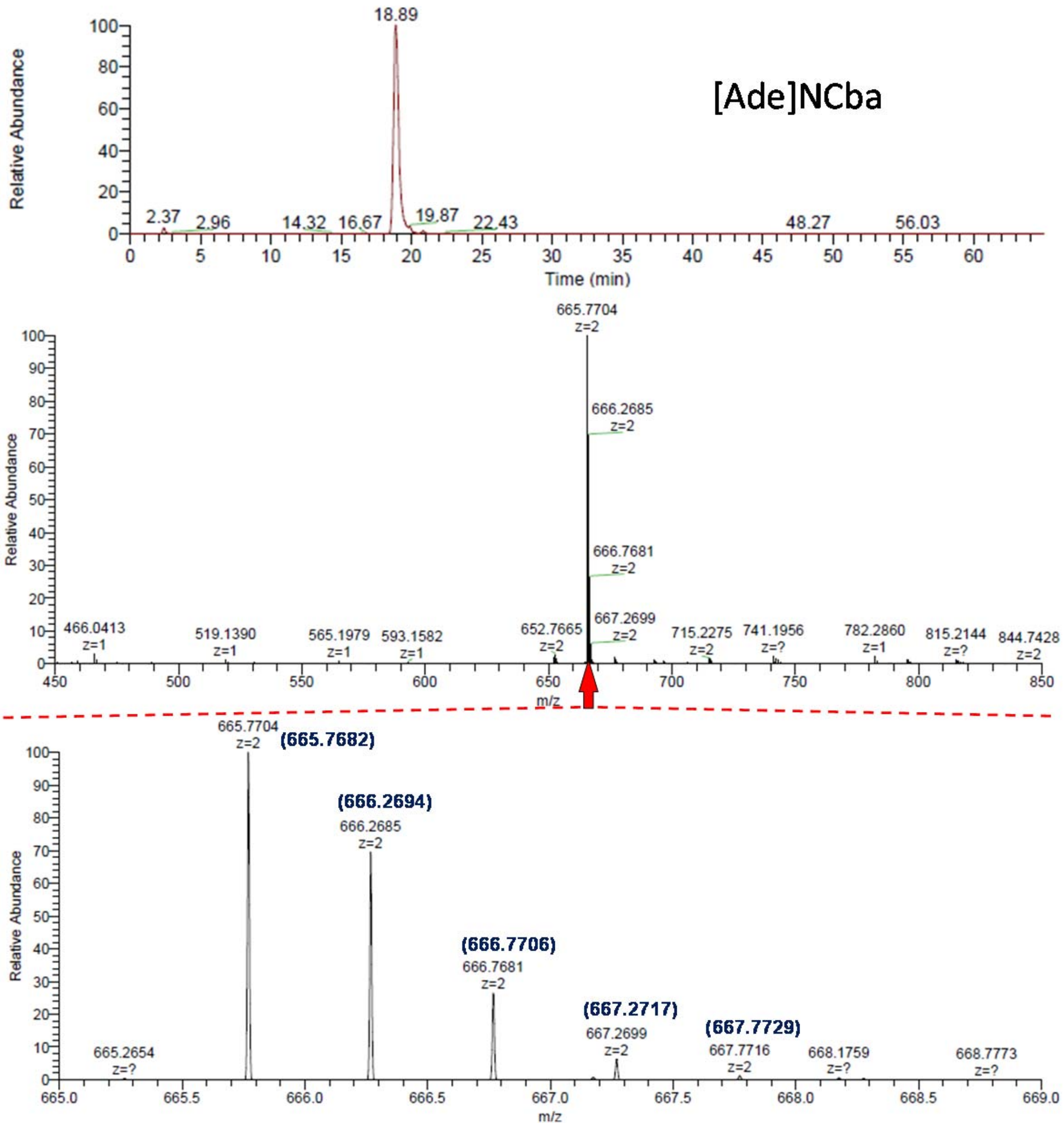


**Supplementary Figure S12: Mass spectrometric analysis of corrinoids extracted from *D. mccartyi* 195 strain grown on $H_2$/*c*DCE with 52 nM [Ade]NCba.** Arrows indicate magnifications of mass spectra. Numbers in brackets represent theoretical mass values of corrinoids to according calculated isotope pattern (see Supplementary Table S1 and S2). [Ade]NCba - adeninyl-norcobamide.

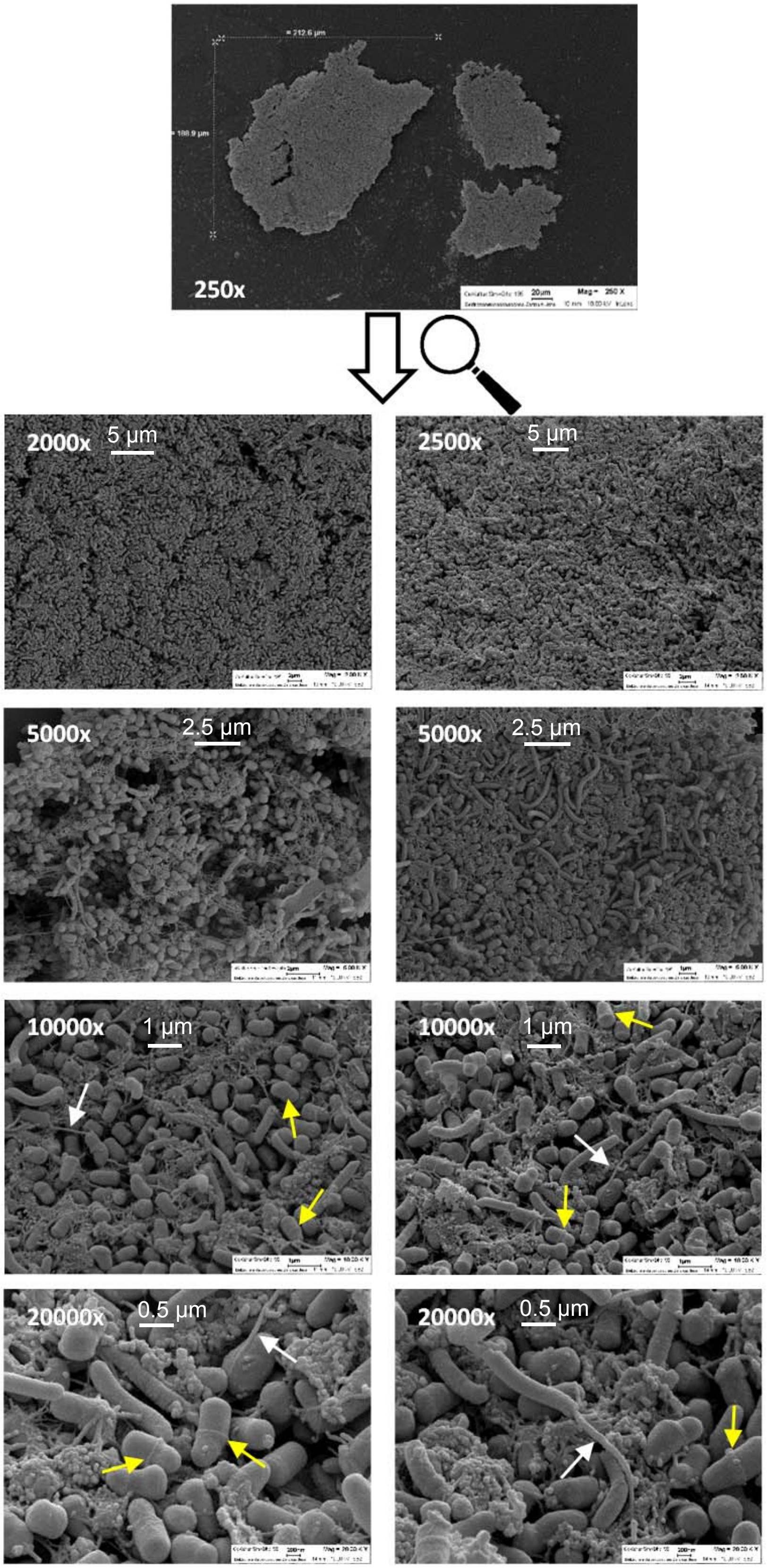


**Supplementary Figure S13: A detailed view and different magnifications of an aggregate of *S. multivorans* and *D. mccartyi* strain 195.** White arrows indicate flagella and yellow arrows indicate ring-shaped septum. Micrographs were taken from different parts of the aggregate. Primary magnifications are given in upper left corner of each micrograph.

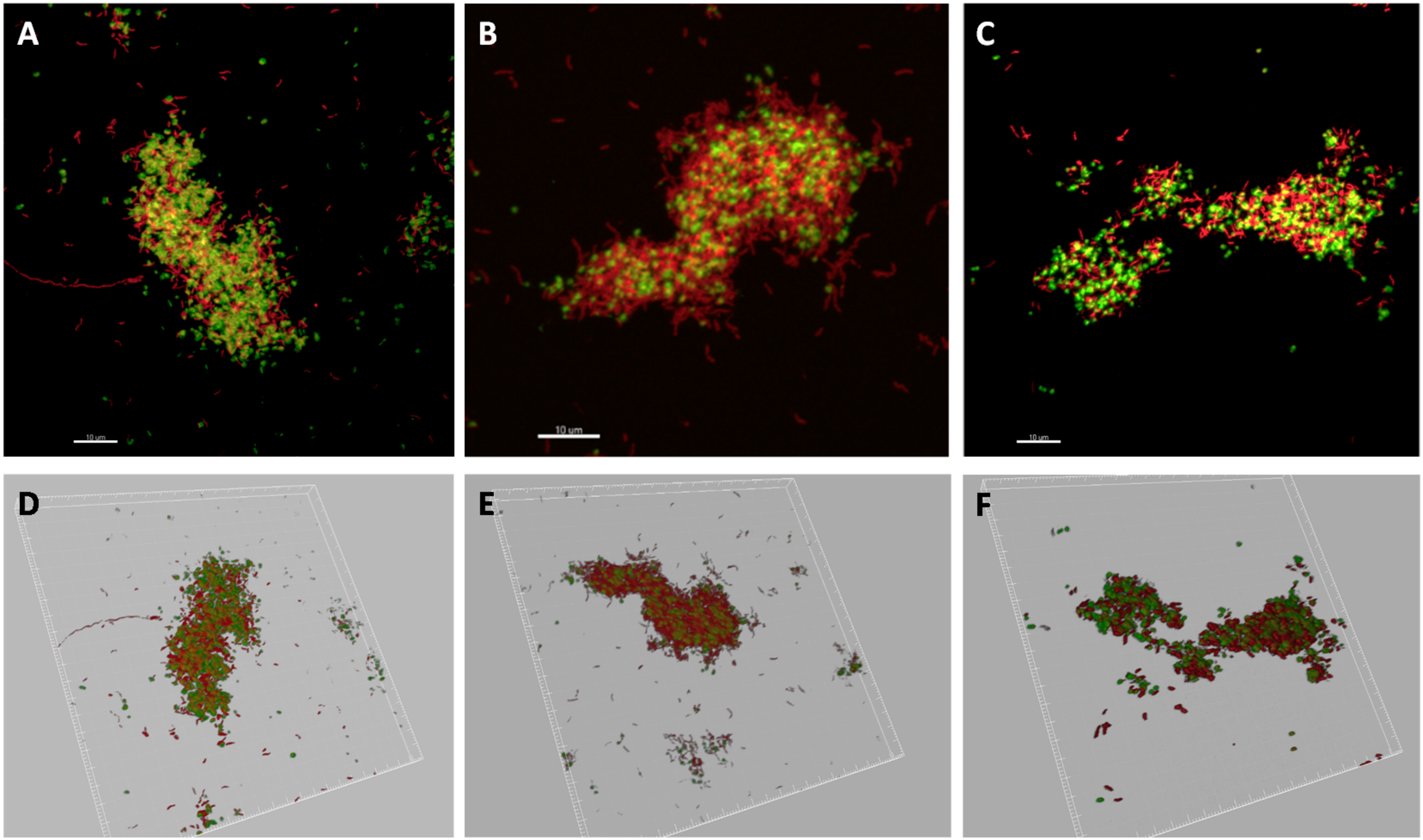


**Supplementary Figure S14: FISH stained aggregates of co-cultures of *S. multivorans* and *D. mccartyi*strain BTF08 (A, D) and 195 (B, C, E, F) and corresponding 3-dimensional imaging (D-F).** red: *S. multivorans*, green: *D. mccartyi.*

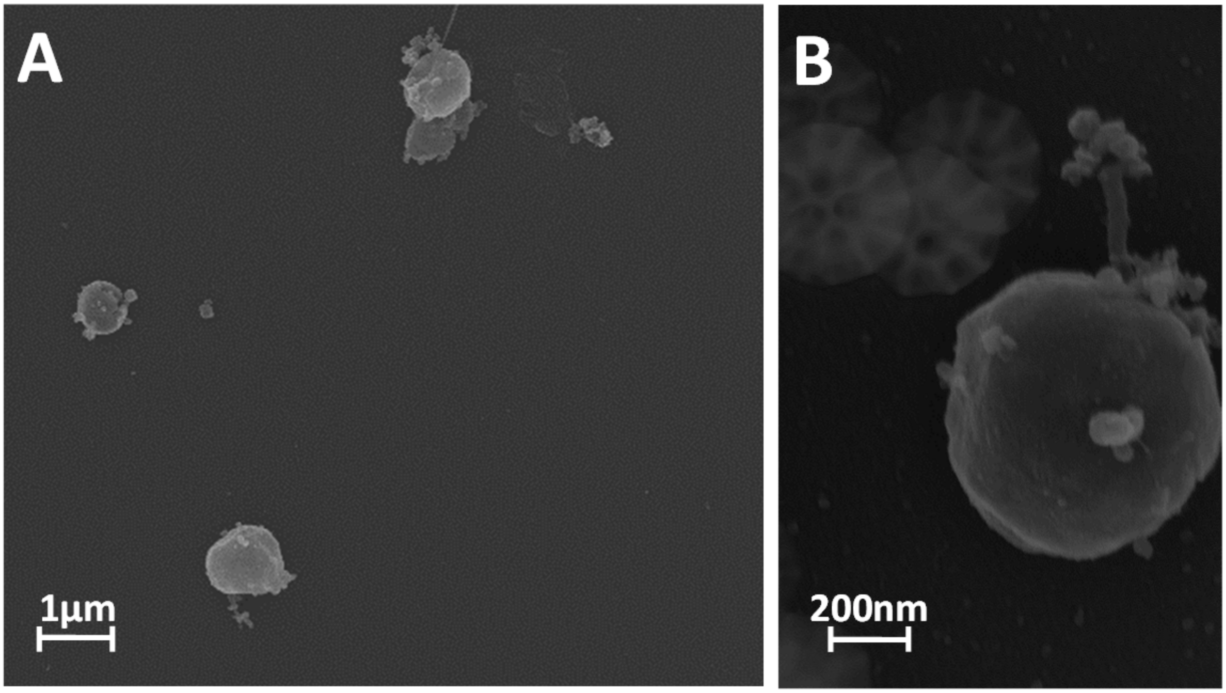


**Supplementary Figure S15:** Cell morphology of *D. mccartyi* strain BTF08 in pure culture.

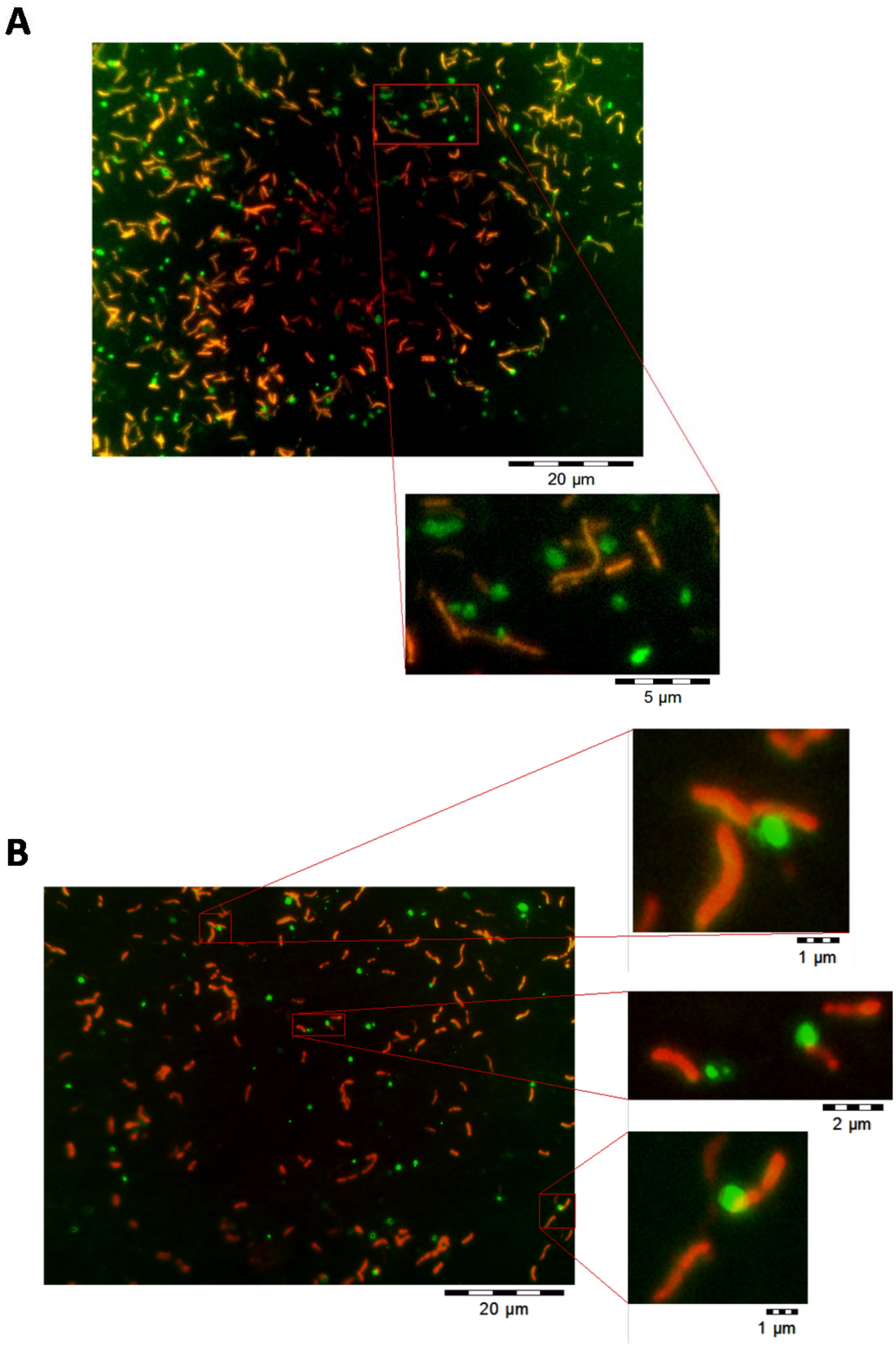


**Supplementary Figure S16: FISH stained co-cultures of *S. multivorans* and *D. mccartyi* strain BTF08 (A) and 195 (B).** Red: *S. multivorans*, green: *D. mccartyi.*

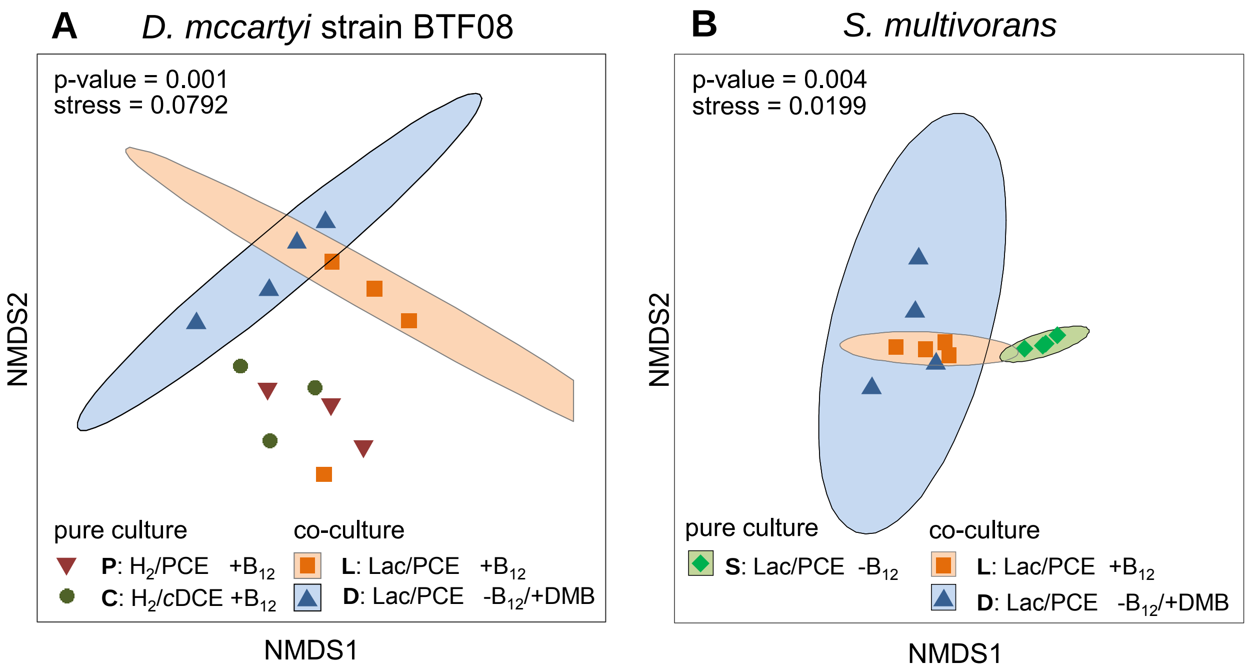


**Supplementary Figure S17: nMDS-analysis of all protein abundances of *D. mccartyi* strain BTF08 (A) and *S. multivorans* (B) cells cultivated under different growth conditions.** Growth conditions are described in Supplementary table 1. For samples C and P only three replicates were available; therefore no confidence interval could be calculated. Stress values indicate that the ordination represents the data very well.

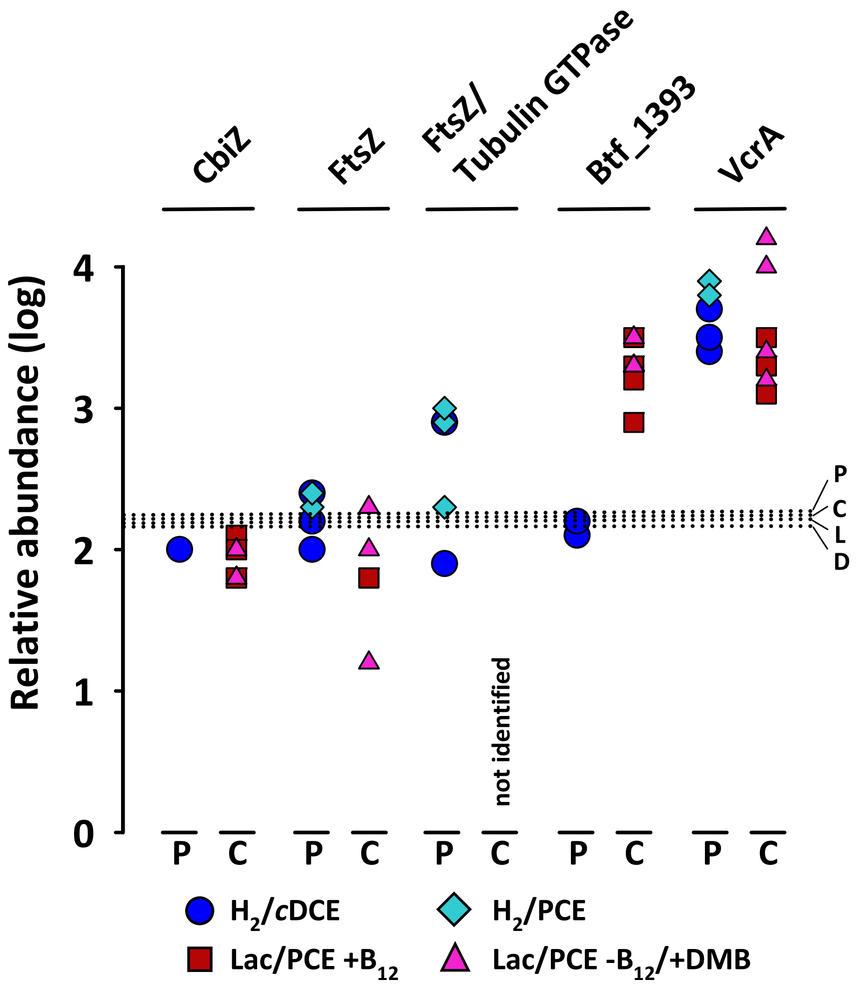


**Supplementary Figure S18: Protein abundances of reductive dehalogenases and a set of significant indicator proteins of *Dhc* BTF08 in pure and co-cultures.** Dots represent lg abundance values from each replicate of the cultures as displayed in figure 5 as average values (pure culture *Dhc* BTF08: $H_2$/*c*DCE [blue circle, N=3], $H_2$/PCE [turquoise diamond, N=3]; co-culture *Sm*/BTF08: Lac/PCE +$B_{12}$ [red square, N=4], Lac/PCE -$B_{12}$/+DMB [pink rectangle, N=4]). Median values of all proteins detected within the different conditions are indicated by dotted lines (C: 2.30, P: 2.33, L: 2.28, D: 2.25). Abundance values of the replicates are given in Supplementary Table S4. CbiZ - adenosylcobinamide amidohydrolase (btf_610), P: *D. mccartyi* BTF08 pure culture, C: *Sm*/BTF08 co-culture, FtsZ - cell division protein (btf_0595), FtsZ/Tubulin GTPase (btf_0551), VcrA - vinyl chloride reductive dehalogenase (btf_1407), btf_1393 - reductive dehalogenase homolog to 11a5_1355 of *D. mccartyi* 11a5.

**Supplementary Table S3: Protein abundances (lg) of molybdopterin-containing oxidoreductases in *S. multivorans* cultivated in co-culture with *D. mccartyi* strain BTF08 and pure culture under different conditions.** Color code represents protein abundances (red: high, green: low). Not detected proteins are colored light grey. $\overline{x}$ - mean values of replicates.

| | | **Co-culture *Sm*/BTF08** | | | | | | | | **Pure culture *Sm*** | | | | $\overline{x}$ | | |
|---|---|---|---|---|---|---|---|---|---|---|---|---|---|---|---|---|
| | | Lac/PCE +$B_{12}$ [L] | | | | Lac/PCE -$B_{12}$/+DMB [D] | | | | Lac/PCE [S] | | | | | | |
| **Accession number** | **Description** | 1 | 2 | 3 | 4 | 1 | 2 | 3 | 4 | 1 | 2 | 3 | 4 | **[L]** | **[D]** | **[S]** |
| **SMUL_0079** | cytoplasmic formate dehydrogenase | 2.0 | 2.3 | 2.4 | 2.5 | 1.9 | 2.2 | 1.2 | 2.1 | 1.2 | 1.5 | 1.5 | 2.0 | 2.3 | 1.9 | 1.6 |
| **SMUL 0273** | molybdopterin oxidoreductase, chain B | | | | | | | | | 2.3 | 2.2 | 2.1 | 1.9 | - | - | - |
| **SMUL_0274** | molybdopterin oxidoreductase, chain A | | | | 1.6 | | | | | | 0.9 | | | - | - | - |
| **SMUL_0342** | polysulfide reductase, subunit A | 2.7 | 2.1 | 2.6 | 2.8 | 2.8 | 2.6 | 2.8 | 2.2 | | | | | 2.5 | 2.6 | - |
| **SMUL_0343** | polysulfide reductase, subunit B | 2.7 | 2.9 | 2.1 | 2.5 | 2.8 | 2.0 | 2.7 | 1.8 | | 0.8 | 0.4 | | 2.5 | 2.3 | 0.6 |
| **SMUL_0344** | polysulfide reductase, subunit c | | | | | | | | | | | | | - | - | - |
| **SMUL_0346** | polysulfide reductase-like protein, subunit A | | | | | | | 1.2 | | | | | | - | - | - |
| **SMUL_0347** | polysulfide reductase-like protein, subunit B | | | | | | | | | | | | | - | - | - |
| **SMUL_0348** | polysulfide reductase-like protein, subunit C | | | | | | | | | | | | | - | - | - |
| **SMUL_0500** | trimethylamine-N-oxide reductase-like protein | 3.2 | 4.1 | 3.5 | 3.8 | 3.7 | 3.7 | 3.6 | 3.4 | | | | | 3.6 | 3.6 | - |
| **SMUL_0501** | cytochrome c-type protein | 3.4 | 2.6 | 3.0 | 2.9 | 3.0 | 2.3 | 2.6 | 3.0 | | | | | 2.7 | 3.0 | 1.6 |
| **SMUL_0912** | nitrate reductase, NapA | 4.3 | 4.5 | 4.5 | 4.4 | 3.8 | 4.7 | 4.1 | 4.5 | 1.9 | 1.9 | 2.0 | 2.0 | 4.4 | 4.3 | 2.0 |
| **SMUL_0913** | nitrate reductase, NapG | | | | | 2.5 | | 2.3 | | | | | | - | 2.4 | - |
| **SMUL_0914** | nitrate reductase, NapH | | | | | 1.9 | | 1.9 | | | | | | - | 1.9 | - |
| **SMUL_0915** | nitrate reductase, NapB | 3.3 | 3.3 | 2.7 | 2.8 | 3.4 | 3.3 | 2.9 | 3.3 | 1.7 | 1.8 | 1.8 | | 3.0 | 3.2 | 1.8 |
| **SMUL_0950** | molybdopterin oxidoreductase chain A | 2.1 | 2.2 | 1.5 | | | 1.7 | 2.0 | 1.6 | | | | | 1.9 | 1.7 | - |
| **SMUL_0951** | molybdopterin oxidoreductase chain A | | | | | | | | | 1.9 | 2.3 | | | - | - | 2.1 |
| **SMUL_0970** | formate dehydrogenase, FdhA | | | | | | | | | | | | | - | - | - |
| **SMUL_0971** | formate dehydrogenase, FdhB | | | | | | | | | | | | | - | - | - |
| **SMUL_0972** | formate dehydrogenase, FdhI | | | | | | | | | | | | | - | - | - |
| **SMUL_1277** | TMAO reductase-like protein | 3.5 | 3.7 | 3.0 | 2.5 | 2.8 | 4.0 | 3.3 | 3.6 | | 1.2 | 1.3 | 1.9 | 3.2 | 3.4 | 1.5 |
| **SMUL_1278** | TMAO reductase-like protein, cytochrome c-type subunit | 3.2 | 3.4 | 1.9 | 1.9 | 2.4 | 3.4 | 3.0 | 3.1 | | | | | 2.6 | 3.0 | - |
| **SMUL_2141** | molybdopterin oxidoreductase | 2.2 | 2.5 | 2.5 | 2.5 | | 2.4 | 2.1 | 2.5 | 1.8 | 2.1 | 2.3 | 2.4 | 2.4 | 2.3 | 2.1 |

| | | | | | | | | | | | | | | | | |
|---|---|---|---|---|---|---|---|---|---|---|---|---|---|---|---|---|
| **SMUL_2312** | dimethylsulfoxide reductase DmsA | 1.8 | 2.0 | 2.1 | 1.7 | 2.2 | 1.8 | 2.0 | 1.8 | 2.7 | 2.7 | 2.8 | 2.6 | 1.9 | 2.0 | 2.7 |
| **SMUL_2313** | dimethylsulfoxide reductase DmsB | | | | | | | | | 2.2 | 2.0 | 2.2 | 2.3 | - | - | 2.2 |
| **SMUL_2314** | dimethylsulfoxide reductase DmsC | | | | | | | | | | | | | - | - | - |
| **SMUL_2568** | molybdopterin oxidoreductase subunit A | 1.6 | 1.6 | 1.4 | 1.3 | 2.0 | 1.5 | 2.5 | 1.8 | | | | | 1.5 | 1.9 | - |
| **SMUL_2569** | molybdopterin oxidoreductase subunit C | | | | | | | | | | | | | - | - | - |
| **SMUL_2570** | molybdopterin oxidoreductase subunit B | | | | | 1.7 | | 1.7 | 1.7 | | | | | 1.7 | - | - |
| **SMUL_2871** | formate dehydrogenase. FdhI | | | | | | | | | | | | | - | - | - |
| **SMUL_2872** | formate dehydrogenase, FdhB | 2.0 | 2.6 | 2.5 | 2.6 | | 2.4 | | | 3.3 | 3.3 | 3.3 | 3.2 | 2.4 | - | 3.3 |
| **SMUL_2873** | formate dehydrogenase, FdhA | 2.8 | 2.6 | 2.9 | 3.0 | 2.5 | 2.7 | 2.2 | 2.5 | 3.5 | 3.5 | 3.5 | 3.4 | 2.8 | 2.4 | 3.5 |
| **SMUL_2899** | formate dehydrogenase, FdhI | | | | | | | | | | | | | - | - | - |
| **SMUL_2900** | formate dehydrogenase, FdhB | 2.9 | 2.9 | 3.0 | 3.1 | 2.4 | 3.0 | 2.7 | 3.0 | 1.9 | 1.9 | 1.9 | 1.9 | 3.0 | 2.8 | 1.9 |
| **SMUL_2901** | formate dehydrogenase, FdhA | 3.7 | 3.6 | 3.8 | 3.9 | 3.2 | 3.7 | 3.3 | 3.6 | 1.5 | 1.7 | 1.9 | 1.9 | 3.7 | 3.5 | 1.7 |
| **SMUL_3029** | Psr/DMSO reductase, chain C | | | | | | | | | | | | | - | - | - |
| **SMUL_3030** | Psr/DMSO reductase, chain B | | | | | | | | | | 1.3 | | | - | - | - |
| **SMUL_3031** | Psr/DMSO reductase, chain A | | 1.3 | | | | | | 1.0 | 1.5 | | | | - | - | - |
| **SMUL_3119** | arsenite oxidase subunit AioB | | | | | | | | | | | | | - | - | - |
| **SMUL_3120** | arsenite oxidase subunit AioB | | | | | | | | | | | | | - | - | - |
| **SMUL_3145** | arsenate reductase. chain C | | | | | | | | | | | | | - | - | - |
| **SMUL_3146** | arsenate reductase. chain A | | | | | | | | | | | | | - | - | - |
| **SMUL_3147** | arsenate reductase. chain B | | | | | | | | | | | | | - | - | - |
| **SMUL_3254** | molybdopterin oxidoreductase, chain C | | | | | | | | | | | | | - | - | - |
| **SMUL_3255** | molybdopterin oxidoreductase, chain B | | | | | | | | | | | | | - | - | - |
| **SMUL_3256** | molybdopterin oxidoreductase, chain A | | | | | | | | | | | | | - | - | - |
| **SMUL_3273** | putative polysulfide reductase, chain C | | | | | 2.5 | | 2.2 | 2.5 | | | | 2.3 | - | 2.3 | - |
| **SMUL_3274** | putative polysulfide reductase, chain B | 2.1 | 2.0 | 2.4 | 2.5 | 3.1 | 1.9 | 2.6 | 1.9 | 1.2 | 1.5 | | | 2.3 | 2.4 | 1.3 |
| **SMUL_3275** | putative polysulfide reductase, chain A | 2.4 | 2.1 | 2.9 | 2.6 | 3.2 | 2.1 | 2.8 | 2.6 | | 1.5 | | | 2.5 | 2.7 | - |
| **SMUL_3281** | putative polysulfide reductase/sulfide dehydrogenase, chain B | | 0.7 | | 1.4 | 1.3 | 1.0 | 1.5 | | 1.6 | 1.9 | 1.8 | 1.7 | 1.0 | 1.3 | 1.8 |
| **SMUL_3282** | putative polysulfide reductase/sulfide dehydrogenase, chain A | 1.7 | | 1.9 | 2.0 | 1.9 | 1.7 | | | | 1.5 | 1.5 | | 1.9 | 1.8 | 1.5 |

**Supplementary Table S4: *D. mccartyi* strain BTF08 protein abundances (lg) of reductive dehalogenases and significant indicator proteins under different cultivation conditions and in pure or co-culture with *S. multivorans*.** $\overline{x}$ - mean value of replicates.

| Accession number | Description | Pure culture *Dhc* BTF08 | | | | | | Co-culture *Sm*/BTF08 | | | | | | | | | | | |
|---|---|---|---|---|---|---|---|---|---|---|---|---|---|---|---|---|---|---|---|
| | | $H_2$/*c*DCE [C] | | | $H_2$/PCE [P] | | | Lac/PCE (+$B_{12}$) [L] | | | | Lac/PCE ($-B_{12}$. +DMB) [D] | | | | $\overline{x}$ | | | |
| | | 1 | 2 | 3 | 1 | 2 | 3 | 1 | 2 | 3 | 4 | 1 | 2 | 3 | 4 | [C] | [P] | [L] | [D] |
| btf_610 | adenosylcobinamide amidohydrolase. CbiZ | | 2.0 | | | | | | 2.1 | 2.0 | 1.8 | 1.8 | | 1.8 | 2.0 | - | - | 2.0 | 1.8 |
| btf_595 | cell division protein FtsZ | 2.0 | 2.4 | 2.2 | 2.3 | 2.4 | 2.4 | | 1.8 | 1.8 | | 2.3 | 1.2 | 2.0 | | 2.2 | 2.4 | 1.8 | 1.6 |
| btf_551 | tubulin/FtsZ GTPase | 1.9 | 2.9 | 2.9 | 2.3 | 2.9 | 3.0 | | | | | | | | | 2.6 | 2.8 | - | - |
| btf_1393 | reductive dehalogenase | 2.1 | 2.2 | | | | 1.9 | 3.5 | 3.3 | 3.2 | 2.9 | 3.4 | 3.3 | 3.3 | 3.3 | 2.2 | - | 3.2 | 3.3 |
| btf_1407 | vinyl chloride reductive dehalogenase. VcrA | 3.4 | 3.4 | 3.5 | 3.9 | 3.9 | 3.8 | 3.5 | 3.3 | 3.3 | 3.1 | 4.2 | 3.2 | 4.0 | 3.4 | 3.5 | 3.9 | 3.3 | 3.5 |
